# EBV Type 1 versus Type 2: A determinant of NK cell anti-tumor activity in Burkitt lymphoma

**DOI:** 10.64898/2026.02.09.704808

**Authors:** Catherine S. Forconi, Lauren Shumate, Zachary Racenet, Viriato M’Bana, Cliff Oduor, Angela Matta, Jeni Melo, Peter O. Oluoch, Boaz Odwar, Juliana Otieno, Terry A. Vik, Festus N’juguna, Ann W. Kinyua, Jeffrey A. Bailey, Christian Münz, Ann M. Moormann

## Abstract

Terminally differentiated CD56^neg^CD16^pos^ NK cells have been described after chronic viral and malaria infections, and in children diagnosed with Burkitt lymphoma (BL). Despite CD56^neg^ NK cells appearing to be poor at direct cytotoxicity, they express high levels of cytotoxic granules (i.e. granzymes, perforin), activation markers, and Fc-γ receptors (CD32 and CD16) that are typically engaged in antibody-dependent cell cytotoxicity (ADCC). In addition, the abundance of CD56^neg^ NK cells strongly correlates with IgG1 and IgG3 plasma levels, which are essential subclasses for ADCC. To determine whether CD56^neg^ NK cells have superior ADCC capacity relative to CD56^dim^ NK cells, we performed ADCC assays using effector cells from pediatric cancer patients and healthy children from malaria endemic regions of Kenya, targeting *in vitro* rituximab-treated commercial and newly established BL cell lines. We found that CD56^neg^ NK cells were indeed capable of *in vitro* ADCC, showing a significant increase of CD107a-mediated degranulation in the presence of rituximab; however, they were not as efficient as CD56^dim^ NK cells. Moreover, we found that the ADCC magnitude was significantly lower against EBV-Type 2 (EBV-T2) BL lines compared to EBV-Type 1 (EBV-T1). EBV-T2 tumor cell lines expressed significantly more lytic viral proteins than EBV-T1, making them more sensitive to direct cytotoxicity. Results from this study highlight the importance of assessing inter-patient variation in NK cell profiles in conjunction with ADCC sensitivity and EBV type within tumor cells when evaluating clinical outcomes for NK-mediated immunotherapies.

**Significance:** EBV type dictates NK cytotoxicity: EBV-T1 BL cells require rituximab for NK killing, while EBV-T2 BL cells are eliminated without antibody assistance, highlighting target-specific immune response to EBV-associated cancers.

## Introduction

Natural Killer (NK) cells play a critical role in antiviral and antitumor immune surveillance(1). Due to their natural cytotoxic receptors (NCRs)(2), NK cells can recognize a wide range of ligands and rapidly eliminate infected or malignant cells. CD56^dim^ NK cells, which express high levels of NCRs, are traditionally known to be the most efficient at both direct and antibody-mediated target cell killing. Our previous work showed that CD56^neg^ NK cells represent a large proportion of peripheral NK cells in malaria-exposed children, with an even more pronounced expansion (up to 68%) observed for pediatric patients diagnosed with Epstein-Barr virus positive (EBV^pos^) Burkitt lymphoma (BL)(3,4). These NK cells, as described by us and others, exhibit poor cytotoxicity in the gold standard K562 direct killing assay(3,5,6). CD56^neg^ NK cells were found in other pediatric (non-Hodgkin lymphoma, nephroblastoma)(3,4) cancer patients, suggesting that their presence is not specific to BL. Moreover, this NK cell phenotype has been identified in adult acute myeloid leukemia(7,8), highlighting the need to better understand their impact on the efficacy of antibody-mediated therapies, such as rituximab which targets CD20-expressing B-cells(9).

BL is an aggressive pediatric B-cell cancer(10) that predominantly affects children aged 5-to-9 years residing in sub-Saharan countries such as Kenya(11). Pediatric BL tumors diagnosed in malaria endemic regions are over 90% EBV^pos^(12), with a third containing EBV-Type2 (EBV-T2) in lieu of EBV-Type1 (EBV-T1)(12). Our team has extensively documented how children living in malaria holoendemic areas become infected with EBV at a very early age (before 2 years)(13), and how chronic *Plasmodium falciparum* malaria co-infections lead to “*immune conditioning”* of CD4^pos^ and CD8^pos^ T-cells to mitigate immunopathology(14–20). This also results in loss of viral control and thus contributes to EBV^pos^ BL tumorigenesis. Yet, how the cytotoxic capacity of NK cells contributes to this oncogenic pathway has not been determined.

Rituximab (Rx) is an IgG1 antibody therapy targeting CD20(9) to induce B-cell death via complement activation, antibody-dependent cellular cytotoxicity (ADCC), and antibody-dependent phagocytosis(9). Closer examination of CD56^neg^ NK cells revealed that despite their purportedly poor direct cytotoxicity, they nevertheless express high levels of cytotoxic granules, activation markers, and notably, high levels of CD32 and CD16 Fc-γ receptors (FcγRs), suggesting their potential for ADCC(4). Through FcγRs, NK cells preferentially recognize the Fc portion of IgG1 or IgG3 antibodies that are typically involved in ADCC(21,22) and thereby trigger cytotoxic responses. In this study, we assessed the relative *in vitro* ADCC activity of CD56^neg^ and CD56^dim^ NK cells using blood collected from children diagnosed with EBV^pos^ BL and from healthy adults and children with divergent malaria exposure histories. Thus, we tested their ADCC activity against a range of BL target cells that were either EBV-T1 or EBV-T2, in the presence or absence of rituximab.

## Methods

*All reagents used are in supp.table1*

### Participants and Ethical approvals

Children diagnosed with BL or other solid tumor cancers were enrolled at Jaramogi Oginga Odinga Teaching and Referral Hospital (JOOTRH), Kisumu and Moi Teaching and Referral Hospital (MTRH), Eldoret, in Kenya. Healthy controls were recruited from malaria-endemic regions in Western Kenya(23,24). Peripheral blood was collected from pediatric cancer patients before chemotherapy. Written informed consent was obtained from each adult participant and child’s guardian. Ethical approvals were obtained from the Kenya Medical Research Institute (KEMRI) Scientific and Ethical Research Unit, MTRH Institutional Research and Ethics Committee, and the University of Massachusetts Chan Medical School Institutional Review Board and followed the principles of the Declaration of Helsinki.

### Demographic data

Participants were classified based on anti-malarial antibody profiles (suppl. fig. 1A) into non-malaria exposed (NME) and malaria-exposed (ME) healthy controls, or as BL cancer patients (supp.table.2). All participants were EBV^pos^ based on VCA and EBNA1 serology (supp. fig. 1). Sex distribution was similar among healthy children and adults (∼55% male), while BL patients had a higher male prevalence (80%) which is typical for this cancer(11). For the ADCC assays, a subset of 10 pediatric cancer patients and 8 healthy children were analyzed (supp.table.3, supp.fig. 1B). NK subset distributions per participant are shown supp.fig.2. Cancer patients (table.2, median 7.5 years, IQR 7–8.25) were slightly younger than healthy controls (median 10 years, IQR 6–11), but the difference was not statistically significant (*p*=0.39). For the survival analysis, 99 confirmed BL patients were included (supp.table.4): 57 BL patients treated with rituximab and standard combined chemotherapy, and 42 treated with our standard combined chemotherapy alone (participants enrolled pre-approval of rituximab as a treatment option)(25). All 99 BL patients were treated at MTRH under the same standard of supportive care.

### Luminex Assay

Total plasma IgG levels against *Plasmodium falciparum* (*Pf*): AMA1, CSP and HRP2; EBV: VCA, EBNA1, Zebra; cytomegalovirus (CMV): pp28; and bovine serum albumin (BSA) as an internal control were measured using Luminex assay following the xMAP-Cookbook protocol(26). Antigens were covalently coupled to magnetic beads following the xMAP-Cookbook protocol. Briefly, after activation of the Luminex magnetic beads, ∼5μg of each antigen was added per million beads to each magnetic bead region. After 2h incubation in the dark on the rotator, beads were washed, counted with a hemocytometer and resuspended in a storage buffer (composition in the xMAP-Cookbook protocol). Coupled beads were kept at 4°C, in the dark, until use. On the day of the Luminex assay, participant’s plasma (dilution 1:100) was incubated with 500 beads per antigen for 2h in a flat-bottom p96 well plate, on a plate shaker (∼800rpm). Secondary biotinylated anti-human IgG (dilution 1:1000) and streptavidin-PE (dilution 1:1000) were added for 1h each. After a last wash, fluorescence was acquired on FlexMap3D, and results reported as antigen-specific median fluorescence intensity (MFI), with BSA values subtracted to account for non-specific binding. The standard curve, made from a pool of plasma from 10 seropositive individuals, used a 2-fold serial dilution; the negative control was the plasma from a seronegative individual (diluted 1/100). Quality control of the Luminex data included a review of the standard curve, the negative control, and a minimum of 50 coupled beads per well.

### Plasma and PBMC isolation

Venous blood was collected in sodium-heparin vacutainers and processed within 2 hours. Hemograms were measured using Mindray BC-3000. Plasma was separated after centrifugation at 1,000g/10mins, stored at –20°C, and cell pellets were resuspended in an equal volume of 1XPBS. Peripheral blood mononuclear cells (PBMCs) were isolated using Lymphoprep and SepMate tubes. PBMCs were cryopreserved at 5×10L cells/mL in freezing medium (90% heat-inactivated fetal bovine serum (HI-FBS), 10%DMSO) and stored overnight at –80°C in Mr. Frosty™ containers before transferring to liquid nitrogen. Samples were shipped using a vapor shipper to maintain the cold chain.

### Antibody isotyping/subclasses

Human isotyping kits were used according to the manufacturer’s instructions, and samples were analyzed on a BioPlex-200 system.

### Target cells

Standard cell lines Daudi, Akata and Mutu (EBV-T1) and our newly established EBV^pos^ BL cell lines: BL717, BL740 (both EBV-T1) and BL719, BL725 (both EBV-T2)(27) were cultured at 37°C/5%CO₂. Akata, Mutu and our newly established EBV^pos^ BL cell lines BL717, BL740 and BL719, BL725 were cultured in RPMI-1640 with 10% heat-inactivated human serum (HI-HS), 1% HEPES, 1% L-Glutamine, 1% NEAA, 1% sodium-pyruvate, and 1% Pen/Strep whereas Daudi was cultured with ATCC media supplemented with 10% HI-FBS. Cell lines with >85% viability (median 96%, range 89–100%) were used for ADCC assays, starting with 5×10L target cells stained with CFSE following the manufacturer’s protocol, while 2×10L remained unstained for controls. Target cells were incubated 30 minutes at 37°C/5%CO₂ with 10Lμg/mL rituximab or isotype control, then washed, and resuspended in complete media. Complete media is composed of RPMI-1640 with 10% HI-HS, 1% HEPES, 1% L-Glutamine, and 1% Pen/Strep.

### Effector cells

PBMCs were thawed and incubated overnight at 37°C/5%CO₂ in NK media (NK-MACS media supplemented with 5% HI-HS, 1% HEPES, IL-2 (200IU/mL), and IL-15 (5ng/mL), at a concentration of 10×10L cells/mL. The following day, PBMCs with viability >85% (median 90%, range 85–98%) were used for ADCC assays.

### ADCC and Flow Cytometry

In a U-bottom 96-well plate, 200k PBMCs were seeded per well. CFSE-labeled and opsonized target cells were added to achieve target:effector (T:E)-ratios of 1:1, 1:3, and 1:9. Each well received a mixture of TAPI-1 (10LμM) and anti-CD107a-PE-Cy5 antibody. To demonstrate technical replicate consistency, we performed the assay in triplicates with Daudi (supp.fig.3) supporting the use of singlets for the other targets. Plates were incubated at 37°C/5%CO₂ for 24h. Cells were washed, resuspended in Zombie-NIR (1/1000) 20mins at room temperature; washed again and resuspended in the antibody-cocktail: CD3-BV421; CD19-PB; CD16-BV650; CD15-PE; CD56-PE-Cy7; CD20-APC; CD14-AF700, for 30mins/4°C. After washing, cells were resuspended in 4%-paraformaldehyde buffer (PBS/2%HI-HS) and passed through 70μm-filters prior to the flow cytometer. Data was acquired on an Aurora using SpectroFlo® software, with compensation controls. Data quality and compensation matrix adjustments were performed using SpectroFlo®. Analysis was done using FlowJo (gating strategy supp.fig.4).

### Clustering analysis using OMIQ

Starting with the PeacoQC algorithm to exclude aberrant events, we ran a FlowSOM clustering on QC-passed live lymphocytes using all the lineage markers. Selecting NK cells positive clusters only, we performed another FlowSOM with CD16, CD56, and CD107a markers to further characterize NK cell subsets (manual gating versus FlowSOM, supp.fig.5). EdgeR was used to compare the abundance of NK subsets between isotype and rituximab conditions, with both *p*-values (≤0.05) and false discovery rate (FDR) scores used to interpret the results.

### Library preparation and RNA sequencing

Total RNA was extracted from BL cell lines using the NEB-RNA extraction kit. Starting with 1–5µg total RNA, we enriched mRNA molecules using oligo-dT magnetic beads to capture polyA-tail mRNA molecules. We then prepared strand-specific RNA-seq libraries following the NEB preparation kit. Final library concentration and fragment size were confirmed with a Qubit-4 fluorometer and fragment analyzer, respectively. These libraries were sequenced with paired-end reads (2×75bp) on the NextSeq-550 system.

### Read Alignment and Gene expression analysis

Sequenced reads were first checked for quality using FastQC(28) and sorted based on the unique sample indexes identified by Novobarcode (Novocraft-Technologies). Residual Illumina 3′-end adapter sequences introduced during the cDNA synthesis were trimmed using cutadapt(29). After residual adapter removal, the reads were mapped to a concatenated human (GRCh38) and EBV (NC_007605 and NC_009334) reference genomes using STAR (Spliced Transcripts Alignment to a Reference) with default parameters. We used RSEM (RNA-Seq by Expectation Maximization) to quantify gene expression and obtain transcript abundance estimates, generating strand specific expected read counts and gene-level count matrices for each cell line. Statistical analyses were performed in R (version 4.3; https://www.R-project.org/). We conducted differential gene expression (DGE) analysis between EBV-T1 and -T2 cell lines using the edgeR package(30), which applies a trimmed mean of M-values (TMM) normalization and a negative binomial modeling approach(31). We excluded any genes with fewer than 5 counts per million across all samples. To account for multiple testing, we applied the Benjamini–Hochberg procedure, considering genes statistically significant if they had an adjusted *p*-value<0.05, FDR<0.05, and a log₂ fold change of >2 or <−2.

### EBV load and EBV typing

EBV load was measured by quantitative Polymerase Chain Reaction (qPCR), as previously described from 50μl of whole blood(32). DNA was extracted from whole blood cell pellets that were resuspended in 1xPBS using DNA extraction kit and following the manufacturer’s instructions with the addition of 10mM DTT and >8 units/mL of Proteinase K then stored at −20°C until use. EBV copies per μg DNA were normalized using copies of EBV per 2 copies of beta actin. Similarly, we used qPCR to differentiate EBV-T1 from -T2. A multiplex PCR reaction using type-specific primers by targeting sequence polymorphisms of EBNA2 were used to identify either the -T1 or -T2 sequence variants. Primer and probe sequences can be found in (Supp.table.1).

### Statistical Analysis

Due to non-normal data distributions (D’Agostino & Pearson test), non-parametric tests were used. For children, antibody isotype and subclass differences were assessed using Kruskal-Wallis with Dunn’s correction; adult comparisons used two-tailed Mann-Whitney tests. ADCC assay outcomes were analyzed as percentage lysis. For Akata, BL717, and BL725, and for Mutu, BL740, and BL719, we used two-tailed Wilcoxon paired-tests to compare isotype vs. rituximab-treated conditions. Median and 95% CI are reported; *p*-values<0.05 are shown. To assess the magnitude of ADCC, we defined cell death as background (spontaneous lysis), direct cytotoxicity (DC; isotype minus background), and ADCC (rituximab minus DC). Finally, effector cell activity was assessed via CD107a expression in NK cell subsets and CD3^pos^ cells. For Akata, BL717, and BL725, Kruskal-Wallis with FDR correction (Benjamini et al.) was used. For Mutu, BL740, and BL719 Wilcoxon paired-tests were applied. Survival was analyzed using the Kaplan-Meier method. Censoring occurred for patients who were event-free at “treatment completion” and “last date known alive” for the short- and long-term survival analysis, respectively. The Logrank (Mantel-Cox) test result is indicated.

### Data Availability

Raw sequencing data files from this study are deposited in the NCBI’s database of Genotypes and Phenotypes (dbGaP) with accession number phs001282.v2.p1. All protocols, flow cytometry, and Luminex raw data are available from the corresponding author upon request.

## Results

### Total IgG, IgG1, and IgG3 levels correlate with abundance of CD56^neg^CD16^pos^ NK cells

To assess the antibody landscape across our study groups, we compared total plasma IgG and IgG subclass levels. No significant differences were observed between children based on malaria exposure status or BL diagnosis (fig. 1A-C). However, cumulative malaria exposure for adults was significantly associated with higher levels of total IgG, IgG1, and IgG3 compared to NME adults (fig. 1A-C; *p*=0.0002, *p*=0.0008, and *p*=0.005, respectively) and compared to ME children (fig. 1A-C, *p*<0.0001 for both IgG and IgG3). Other antibody comparisons were made (supp.fig.6A-D) and found to be either non-significant or not relevant to NK cell activity. Consistent with our hypothesis, we found significant positive correlations between CD56^neg^ NK cell abundance and levels of total IgG (fig. 1G, r=0.47, *p*=0.004), IgG1 (fig. 1H, r=0.2, *p*=0.009), and IgG3 (fig. 1I, r=0.49, *p*=0.002). In contrast, no correlations were observed for total IgG2, IgG4, IgA, or IgM levels (supp.fig.6E-H).

**Fig. 1:**
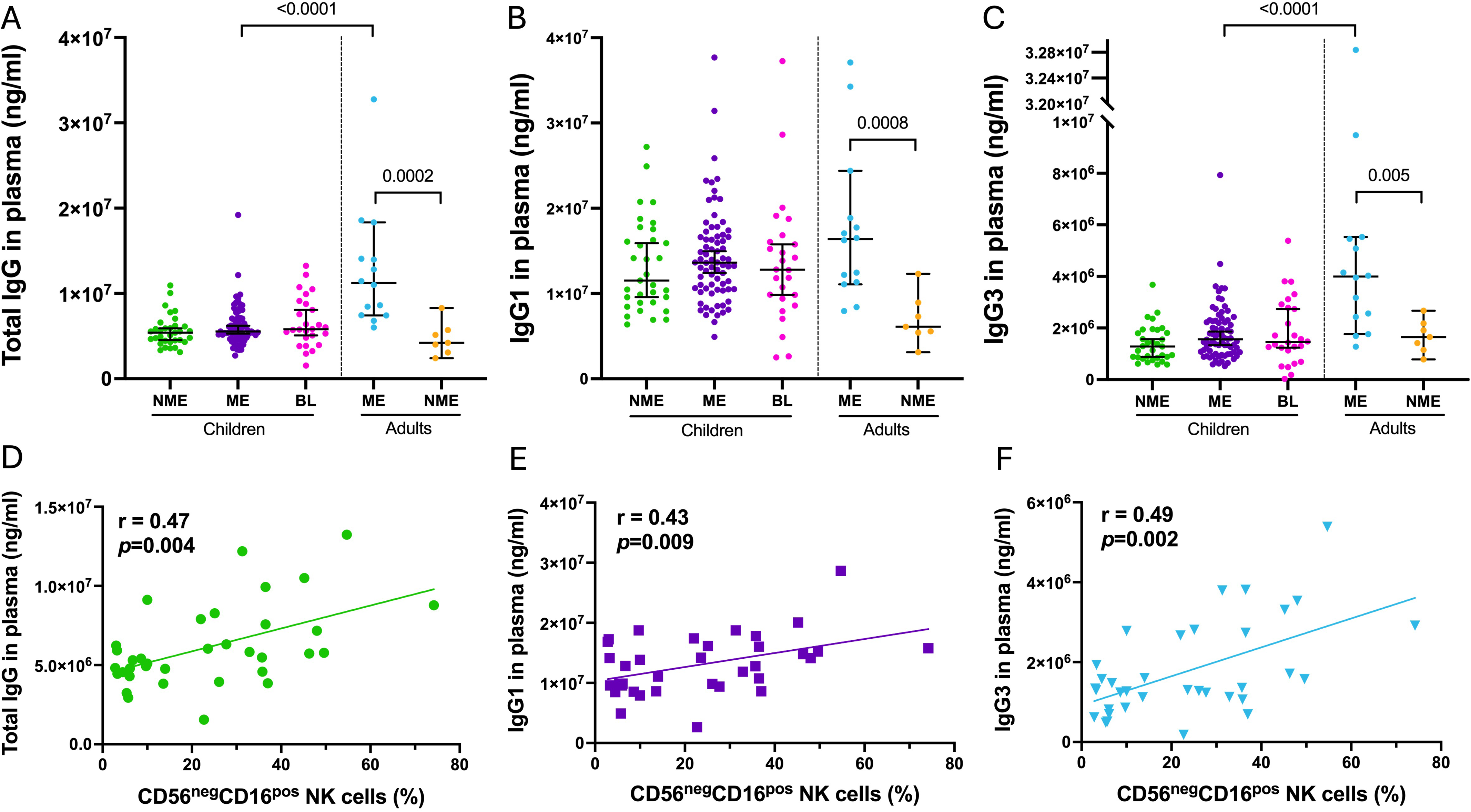
Correlation between total IgG, IgG1 and IgG3 with abundance of CD56^neg^CD16^pos^ NK cells in Kenyan children. Dot plots of plasma level of total IgG **(A)**, IgG1 **(B)** and IgG3 **(C)** from non-malaria exposed (NME, n=33, green) and malaria exposed (ME, n=74, purple) children, BL patients (BL, n=26, pink), ME and NME adults (n=14, blue and n=7, orange, respectively). Median and 95% confidence intervals are represented; no statistical differences were found using Kruskal-Wallis test between the groups of children. Significant *p*-values are indicated following Mann-Whitney test between the two groups of adults and between the ME children and adults. Correlation test assessing relationships between plasma levels of total IgG **(D)**, IgG1 **(E)** and IgG3 **(F)**, and the frequency of peripheral CD56^neg^CD16^pos^ NK cells from total NK cells from all children participants. Plasma levels of total IgG and IgG3 did not follow a normal distribution, whereas IgG1 and the abundance of CD56^neg^CD16^pos^ NK cells did. Therefore, the Spearman test was used for IgG1 and IgG3 while Pearson was used for IgG1 (r and *p*-values indicated on the plots).

### Rituximab enhances lysis of EBV-T1 but not EBV-T2 BL cells

To determine whether the target cell influenced ADCC-mediated lysis induced by rituximab, we compared several BL lines. As expected, rituximab significantly increased lysis at a 1:1 target-to-effector (T:E)-ratio, from 9.7% to 21.5% for Akata-T1 (fig.2A, *p*=0.002), 6% to 16.8% for BL717-T1 (fig.2B, *p*=0.001), and 15.8% to 23.5% for BL725-T2 (fig.2C, *p*=0.001).

**Fig. 2:**
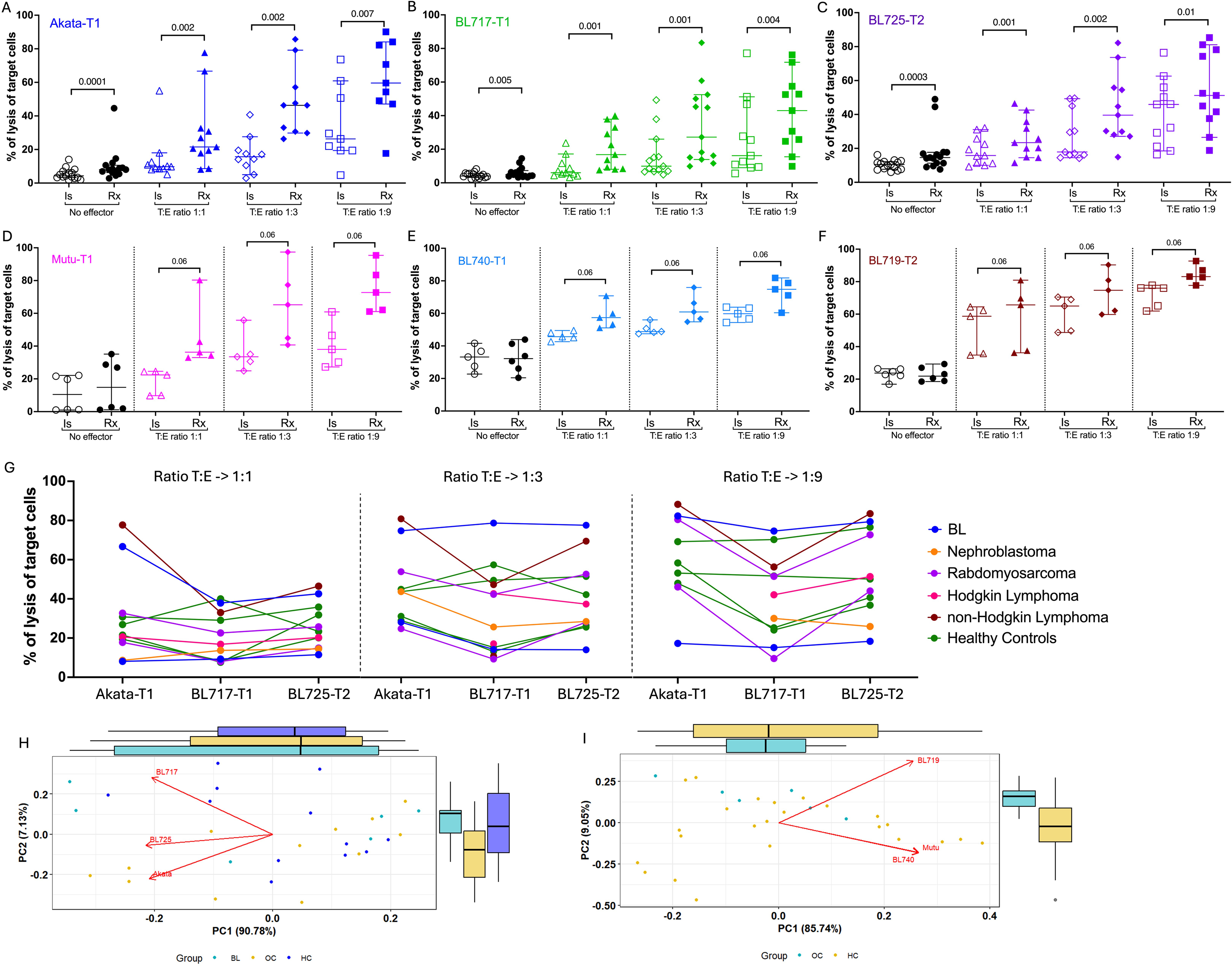
Rituximab enhanced the killing of the target cells regardless of the clinical status of the children. Percentage of lysis of Akata-T1 **(A)**, BL717-T1 **(B)**, BL725-T2 **(C)**, Mutu-T1 **(D)**, BL740-T1 **(E)** and BL719-T2 **(F)**. Data points are represented with median and 95% confidence intervals (n=11 to 13). Isotype condition is shown using open symbols while solid symbols indicate rituximab treatment. The circles represent the percentage of lysis of the target cells in absence of effector cells (background), while triangles, diamond, and squares show the T:E ratio (1:1, 1:3 and 1:9, respectively). Two-tailed paired Wilcoxon test was used for statistical significance, *p*-values are indicated on the plots. Percentage of lysis of the target cells (Akata-T1, BL717-T1 and BL725-T2) highlighting the clinical status of each participant at T:E ratio 1:1, 1:3, and 1:9 **(G)**. Principal component analysis of the percentage of lysis of the two sets of targets: Akata-T1, BL717-T1 and BL725-T2 **(H)** and Mutu-T1, BL740-T1 and BL719-T2 **(I)** in regard to the clinical status of the participant. Acronyms are as follows, non-HL: non-Hodgkin Lymphoma, BL: EBV-associated Burkitt lymphoma, RM: Rhabdomyosarcoma, HC: Healthy control, HL: Hodgkin Lymphoma, T1 and T2 refer to EBV type 1 and type 2, respectively and N: Nephroblastoma.

A progressive increase in lysis was observed with higher effector cell numbers. Interestingly, at a 1:9 (T:E)-ratio, BL725-T2 showed a smaller difference between isotype and rituximab conditions (45.9% vs. 51.2%) compared to >2-fold differences for Akata-T1 (26.3% vs. 59.6%) and BL717-T1 (16.2% vs. 43%). To validate this finding, we replicated this experiment using additional cell lines: Mutu-T1 and BL740-T1, and BL719-T2. Despite variability observed in the second set, rituximab consistently enhanced lysis of the EBV-T1 tumors (fig.2D-E), with median lysis increases of ≥15% compared to isotope controls (Mutu-T1: 38% to 72.7%; BL740-T1: 59.8% to 74.8%). In contrast, lysis of BL719-T2 showed only a 7% increase (76% to 83%) with rituximab treatment (fig.2F). These results suggest that EBV-T2 BL tumors may be more prone to direct cytotoxicity, limiting the contribution of ADCC, while EBV-T1 tumors are more sensitive to antibody-mediated lysis.

### Target cell lysis does not differ between cancer patients and healthy controls

As we previously reported(3,4), we observed a range of CD56^neg^ NK cells (1-41.3%) in both healthy and cancer children (supp.fig.2). No difference in the percentage of target cell lysis was found between cancer patients and healthy controls; however inter-individual lysis differences were likely to be consistent across the different target cells (fig.2G). This finding was further supported by principal component analysis, which revealed no group-based clustering in target cell killing efficiency (fig.2H-I). Importantly, we found no correlation between the percentage of lysis of the rituximab-treated target cells and the percentage of CD56^neg^ NK cells in our study populations (supp.fig.7), suggesting that the mere abundance of CD56^neg^ NK cells did not influence *in-vitro* rituximab-mediated ADCC efficiency.

### Direct cytotoxicity trounces ADCC against EBV-T2 BL tumors

Analyzing the relative contributions of direct cytotoxicity (DC) and ADCC to target cell lysis, we observed that ADCC was more prominent against Akata and BL717-T1 (fig.3A-B, respectively) compared to BL725-T2, which exhibited greater DC, especially at a T:E-ratio of 1:9 (fig.3C). This explains the previously noted minimal difference in lysis between isotype and rituximab conditions for BL725-T2. However, we also observed variability in DC versus ADCC contributions across participants (fig.3D-F). Replication experiments using Mutu-T1 showed greater ADCC at a 1:1 T:E-ratio (supp.fig.8A/D) while BL719-T2 was predominantly killed through DC regardless of the T:E-ratio (supp.fig.8C/F). Interestingly, BL740-T1 also exhibited more DC than ADCC, though to a lesser extent than BL719-T2 (supp.fig.8B/E). Together, these results suggest that rituximab-mediated ADCC acts as an additive cytotoxic mechanism against EBV-T1 tumors but is not the dominant mode of target cell killing for EBV-T2 tumors.

**Fig. 3:**
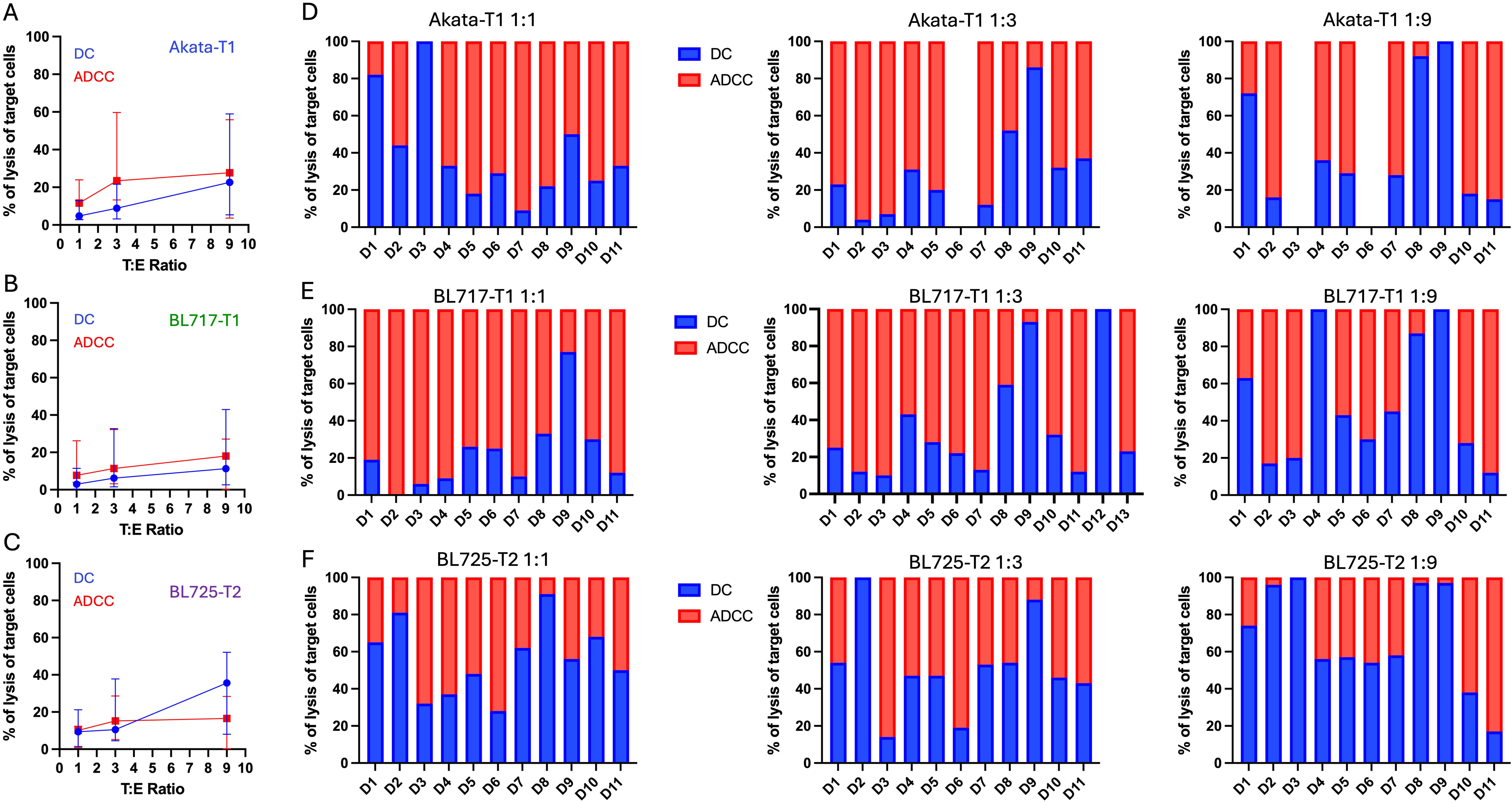
EBV-T1 BL cell lines are preferentially killed via ADCC whereas EBV-T2 BL targets are killed via direct cytotoxicity. Range of percentage of lysis of Akata-T1 **(A)**, BL717-T1 **(B)**, and BL725-T2 **(C)** by DC (in blue) or ADCC (in red) across three different T:E ratios. The symbol represents the median and the bar the error with 95% confidence interval. The proportions of DC (in blue) and ADCC (in red) killing of Akata-T1 **(D)**, BL717-T1 **(E)**, and BL725-T2 **(F)** are then reported with bar plots for each participant for the three different T:E ratios (D1 to D11 for all conditions except BL717-T1 T:E ratio 1:3 which has donors D1 to D13).

### EBV-T2 BL cells express more EBV lytic genes compared to EBV-T1

To explore transcriptional differences between our new pediatric patient-derived EBV-T1 and -T2 BL cell lines, we performed differential gene expression analysis (fig.4). We identified 105 differentially expressed genes (logLFC>2, FDR<0.05, adj-*p*<0.05), including 13 EBV lytic genes such as *BZLF1* and *BMLF1* (Supp.table.5). These lytic genes were significantly upregulated in EBV-T2 BL cells, with expression increased over 16-fold (logLFC<-5) compared to EBV-T1 (fig.4A). Additionally, several host immune-related genes were differentially expressed. Notably, *IL32,* linked to enhanced NK cell function(33), was significantly upregulated in EBV-T2 BL cells (logLFC<-5, FDR<0.0001). *LGALS1*, gene of galectin-1 protein which can be secreted in tumor micro-environment to influence the immune response(34), was also markedly increased in EBV-T2 (fig.4B, logLFC=-7.8, FDR=6.99e-17, adj-*p*=3.15e-13). Interestingly, no change in NK ligand expression was observed transcriptionally (fig.4C/D) or at the protein level (fig.4E) between the EBV-T1 and -T2 cell lines, suggesting that other ‘kill-me’ signals might be in play, such as cytokines. Importantly, all cell lines expressed CD20 but with various intensity regardless of EBV-type (fig.4E/F). Overall, these findings reveal that EBV-T1 BL cell lines exhibit a more latent viral phenotype and downregulate key immune-stimulatory host genes, potentially aiding immune evasion. In contrast, the more lytic profile of EBV-T2 BL cells may increase their visibility to NK cells, contributing to their heightened susceptibility to direct immune clearance.

**Fig. 4.**
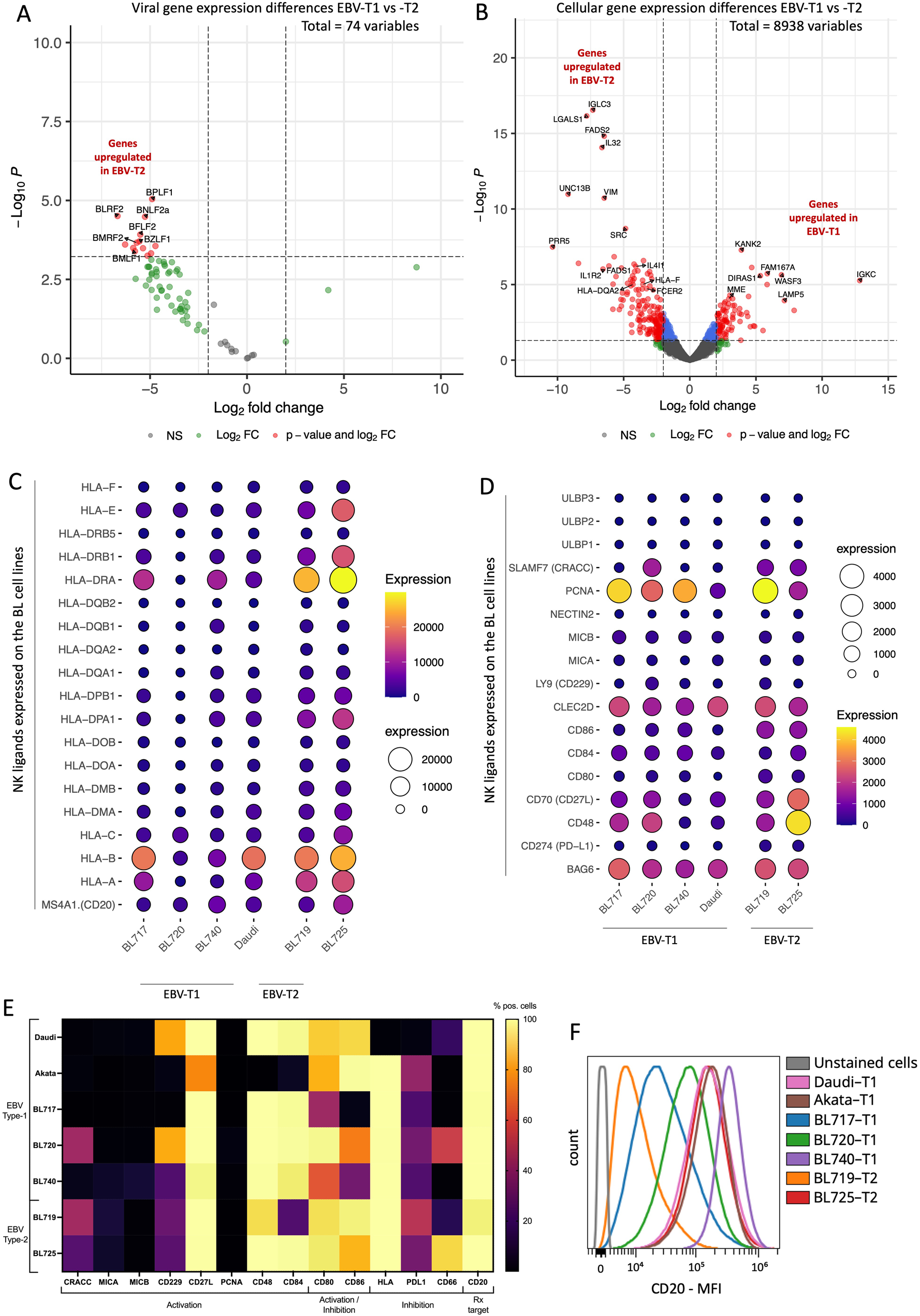
: EBV-T2 BL cell lines expressed higher levels of EBV lytic genes compared to EBV-T1. Volcano plot of differentially expressed viral **(A)** and cellular **(B)** genes between EBV-T1 and EBV-T2 BL cell lines. Each gene tested is plotted by its log2 fold change (logFC) on the x-axis and its −log10 *p*-value on the y-axis. Significantly differentially expressed (DE) genes (logFC > 2, *p*-value < 0.05, FDR < 0.05) are highlighted in red, including 80 downregulated and 25 upregulated genes in EBV-T1 BL cell lines compared to -T2 BL cell lines. Notably, several EBV lytic genes (such as, *BLRF2*, *BZLF1*, *BMLF1*) are downregulated in EBV-T1 BL cell lines, suggesting a potential difference in viral gene expression regulation between these two EBV subtypes. Balloon plots of HLA along with CD20 expression **(C)** and known NK ligands **(D)** at transcriptional level across EBV-T1 and EBV-T2 cell lines. **(E)** Heatmap of selected NK ligands assessed by flow cytometry along with CD20 on EBV-T1 and -T2 cell lines. The color scale indicates the percentage of positive cells, yellow being 100%. **(F)** Histogram showing the CD20 intensity expression (MFI) between the different BL cell lines, the grey being the unstained cells.

### CD56^neg^ NK cells can perform rituximab-driven ADCC

To evaluate effector cell activity, we measured CD107a, marker of NK cell degranulation and cytotoxic function(35). Rituximab significantly increased the frequency of CD56^dim^CD107a^pos^ NK cells compared to isotype and background controls when targeting EBV-T1 cells (Akata and BL717-T1), independent of the T:E-ratio (fig.5A-B, q-value<0.001 and q-value<0.02). In contrast, the effect was weaker against BL725-T2 cells, with significance observed only at a T:E-ratio of 1:9 (fig.5C, q-value<0.02). Interestingly, CD56^neg^CD107a^pos^ NK cells showed similar patterns (fig.5D-F) revealing that CD56^neg^ NK cells can mediate rituximab-driven ADCC against both EBV-T1 and -T2 targets; and because of the similarity between isotype and rituximab conditions, CD56^neg^ NK cells may also contribute to direct cytotoxicity against newly established BL cell lines. However, not all participants had CD56^neg^ NK cell responses, even in the presence of rituximab, whereas CD56^dim^ NK cells consistently responded. Similar trends were observed with Mutu-T1 (supp.fig.9A-C), BL740-T1 (supp.fig.9D-F), and BL719-T2 (supp.fig.9G-I): rituximab broadly enhanced CD56^dim^CD107a^pos^ NK cell activity, while CD56^neg^ responses were limited to fewer donors. These findings suggest that variability in CD56^neg^ NK cell function may cause differences in patient responses to rituximab therapy. Importantly, we also found that the frequency of responding CD56^neg^CD107a^pos^ cells strongly correlated with the percentage of responding CD56^dim^CD107a^pos^ cells, regardless of the target cell used (fig.5G-I). This indicates that CD56^neg^ NK cells complement rather than replace ADCC activity mediated by CD56^dim^ NK cells.

**Fig. 5:**
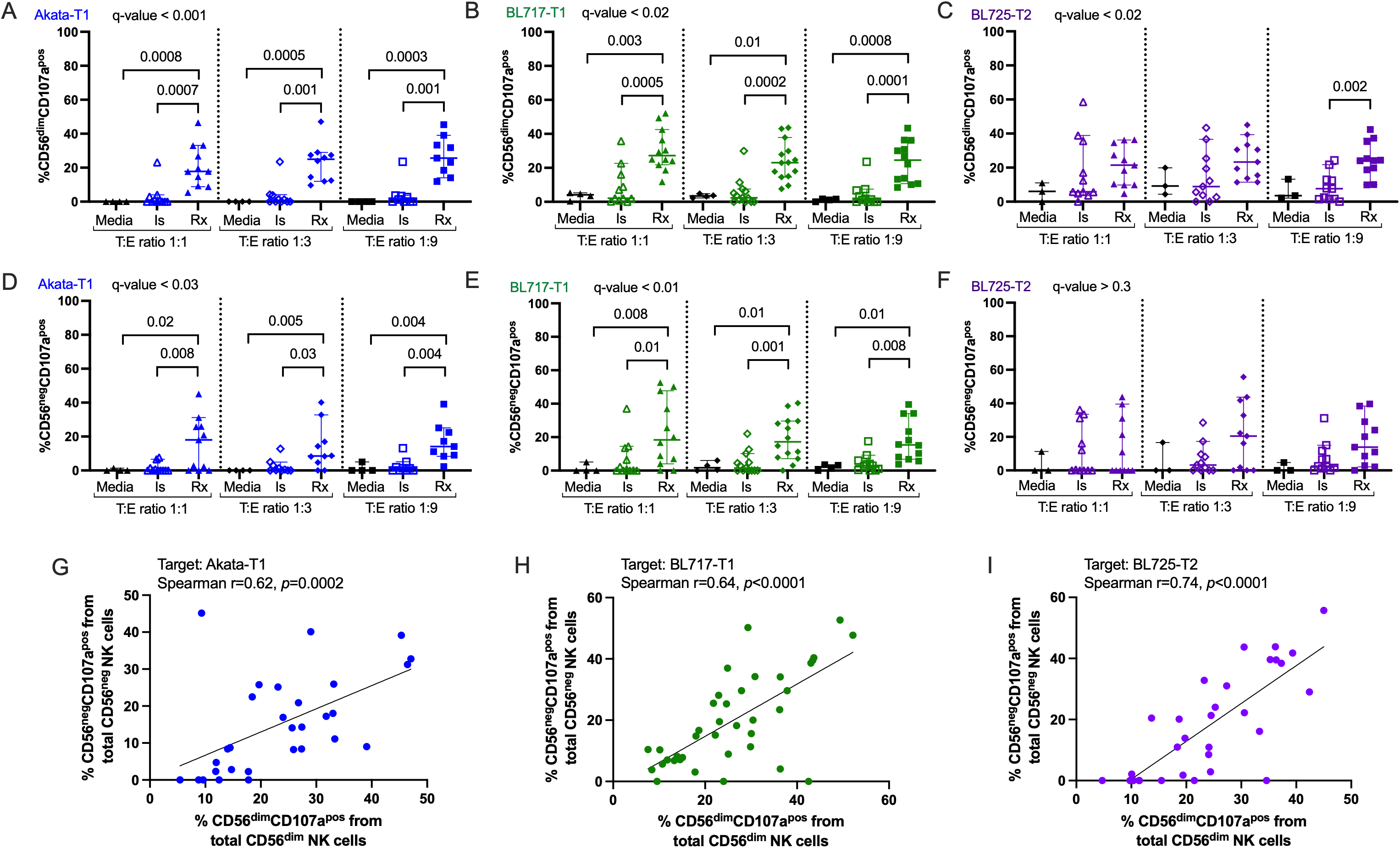
More CD56^dim^ and CD56^neg^ NK cells degranulate in presence of rituximab against EBV-T1 BL cell lines. Percentage of CD56^dim^CD107a^pos^ cells from total CD56^dim^ NK cells after co-culture with Akata-T1 **(A)**, BL717-T1 **(B)** and BL725-T2 **(C)**, in presence of isotype control (empty symbols), rituximab (full symbol) or nothing (black symbols) using manual gating. Percentage of CD56^neg^CD107a^pos^ cells from total CD56^neg^ NK cells after co-culture with Akata-T1 **(D)**, BL717-T1 **(E)** and BL725-T2 **(F)**, in presence of isotype control (open symbols), rituximab (solid symbol) or nothing (black symbols) using manual gating. All the data are represented with median and 95% confidence intervals. Kruskal-Wallis statistical test was performed, and the false discovery rate (FDR) was used to correct for multiple comparisons. Both *p*-values and *q*-values (adjusted *p*-values based on the FDR) are indicated on the plots. Because of the non-normal distribution of the data, the Spearman test was used to assess relationships between the percentages of CD56^dim^CD107a^pos^ and CD56^neg^CD107a^pos^ NK cells, in presence of the target Akata-T1 **(G)**, BL717-T1 **(H)** and B:725-T2 **(I)** (r and *p*-values indicated on the plots).

### CD56^neg^ NK cells exhibit reduced ADCC compared to CD56^dim^ NK cells

To determine whether CD56^neg^ NK cells exhibit greater ADCC activity than CD56^dim^ NK cells, based on their higher FcγRs expression we compared their cytotoxicity against various BL target cells. CD56^dim^CD107a^pos^ NK cells were significantly more abundant than CD56^neg^CD107a^pos^ NK cells under isotype conditions, and this pattern persisted following rituximab treatment (fig.6A-C, q-value<0.01). However, the intensity of degranulation was significantly lower in CD56^neg^CD107a^pos^ NK cells compared to CD56^dim^CD107a^pos^ NK cells when co-cultured with Akata-T1 or BL717-T1 targets (fig.6D-E, q-value<0.01, *p*=0.002 and *p*=0.0001, respectively). This difference was not observed with the BL725-T2 target cells (fig.6F, q-value<0.01). Notably, CD56^dim^CD107a^pos^ NK cells exhibited significantly increased degranulation in the presence of rituximab compared to isotype controls for both Akata-T1 and BL717-T1 (fig.6D-E, q-value<0.01, *p*=0.0008 and *p*<0.0001, respectively). However, with BL725-T2, no statistical difference in degranulation intensity was observed between rituximab and isotype conditions (fig.6F, q-value<0.01). Further analysis revealed that, under isotype conditions, CD56^dim^CD107a^pos^ NK cells degranulated significantly less when co-cultured with Akata-T1 and BL717-T1 compared to BL725-T2 (fig.6G, q-value=0.01, *p*=0.002 for both). These trends were consistent across a second set of target cell lines (i.e. Mutu-T1, 740-T1, and BL719-T2 (supp.fig.9J-O). Altogether, these results indicate that overall CD56^neg^ NK cells have a reduced capacity to perform ADCC compared to CD56^dim^CD107a^pos^ NK cells, when targeting an EBV-T1 BL cell.

**Fig. 6:**
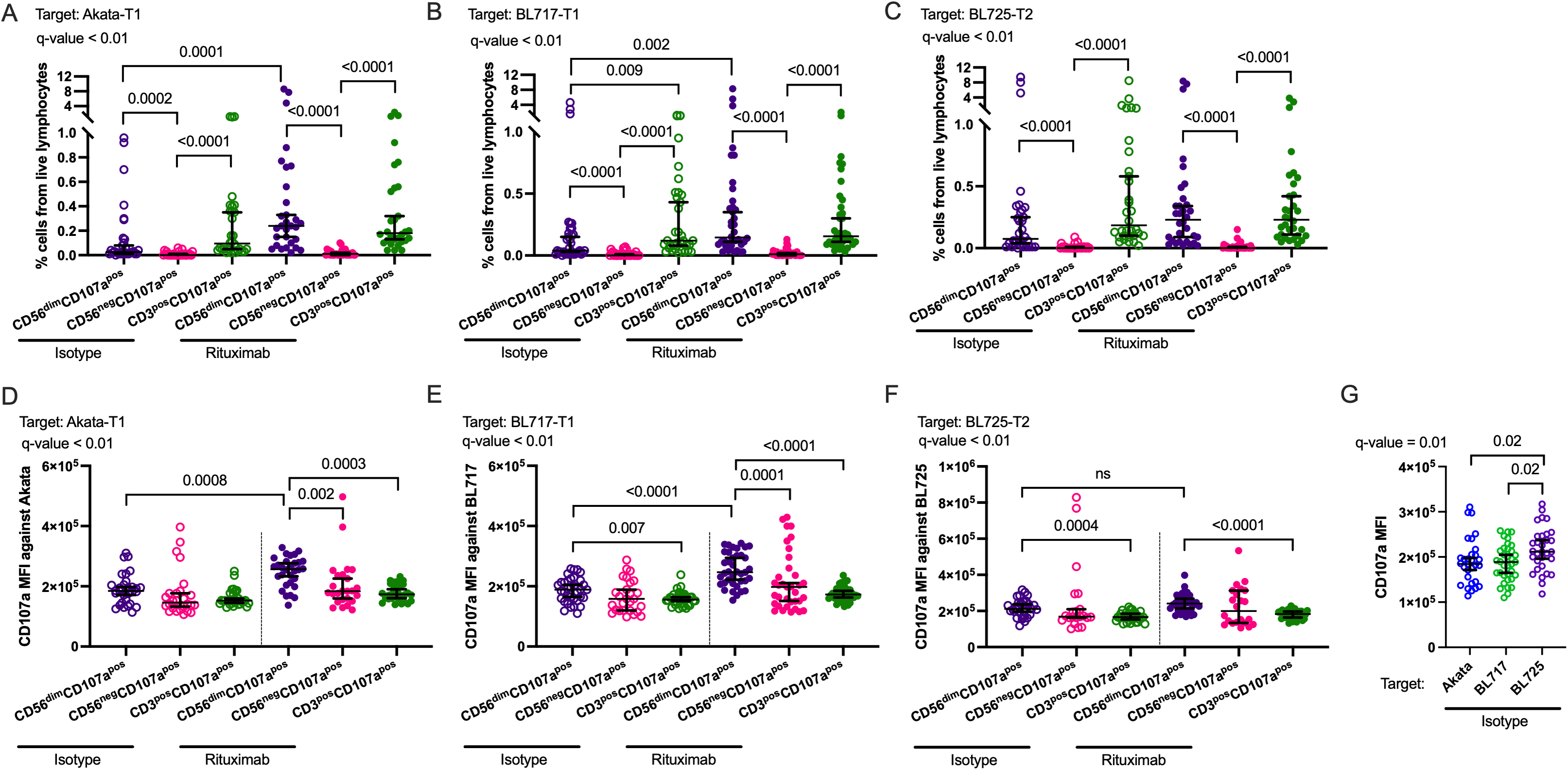
CD56^dim^ NK cells degranulated significantly more than the CD56^neg^ NK cells in presence of rituximab against EBV-T1 BL cell lines. Percentage of CD56^dim^CD107a^pos^ (purple), CD56^neg^CD107a^pos^ (pink), and CD3^pos^CD107a^pos^ (green) cells from total live lymphocytes after co-culture with Akata-T1 **(A)**, BL717-T1 **(B)** and BL725-T2 **(C)**, in presence of isotype (empty symbols) or rituximab (full symbol). CD107a median fluorescence intensity (MFI) of CD56^dim^CD107a^pos^ (purple), CD56^neg^CD107a^pos^ (pink), and CD3^pos^CD107a^pos^ (green) cells after co-culture with Akata-T1 **(D)**, BL717-T1 **(E)** and BL725-T2 **(F)**, in presence of isotype (empty symbols) or rituximab (full symbol). Comparison of the CD107a MFI of the CD56^dim^CD107a^pos^ cells between isotype conditions against Akata-T1 (blue), BL717-T1 (green) and BL725-T2 (purple). All the data are represented with median and 95% confidence intervals. Kruskal-Wallis statistical test was performed, and the false discovery rate (FDR) was used to correct for multiple comparisons. Both *p*-values and *q*-values (adjusted *p*-values based on the FDR) are indicated on the plots.

### Clustering analysis confirms superior ADCC efficiency of CD56^dim^ NK cells

To mitigate potential bias associated with traditional 2D-gating strategies, we conducted a semi-supervised clustering analysis using the FlowSOM algorithm. This approach enabled a more granular identification of NK cell subsets. Statistical comparison of cluster abundance between isotype and rituximab-treated samples was performed using EdgeR for each target cell line. Consistent with our 2D-gating results, CD56^dim^ NK subsets (expressing either low or high levels of CD107a) emerged as the most abundant populations engaged by rituximab, when Akata-T1 or BL717-T1 cells were the targets (fig.7A-C and 7D-F, respectively). CD56^neg^CD107a^pos^ NK cells ranked as the second or third most abundant subset in these conditions. In contrast, consistent with earlier findings that lysis of BL725-T2 involves both direct cytotoxicity and ADCC, no NK cell subset showed a statistically significant difference in abundance between isotype and rituximab conditions (FDR<0.05), regardless of the E:T ratio (fig.7G-I). These findings were recapitulated with the additional BL cell lines Mutu-T1, BL740-T1, and BL719-T2 (supp.fig. 10). Together, these results further support that while CD56^neg^ NK cells are capable of performing ADCC, CD56^dim^ NK cells are more efficient effectors mediating rituximab cytotoxicity.

**Fig. 7:**
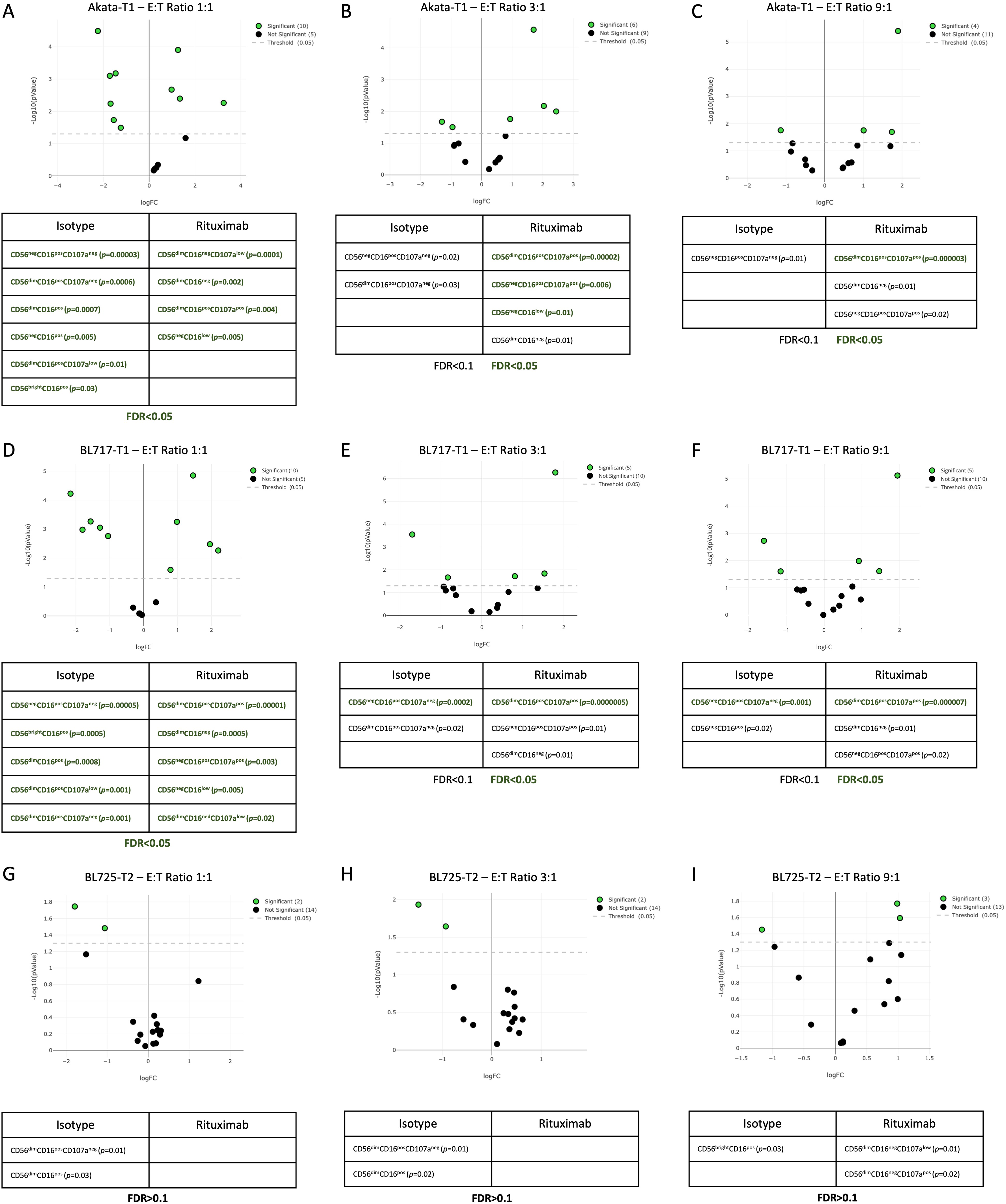
CD56^dim^ NK cells remain superior to ADCC compared to CD56^neg^CD16^pos^ NK cells except against EBV-T2 tumors. EdgeR analysis following a semi-supervised FlowSOM clustering showing, in green, the NK subsets statistically differently abundant between the isotype and the rituximab condition after co-culture with Akata-T1 T:E ratio 1:1 **(A)**, 1:3 **(B)**, 1:9 **(C)**; BL717-T1 T:E ratio 1:1 **(D)**, 1:3 **(E)**, 1:9 **(F)**; and BL725-T2 T:E ratio 1:1 **(G)**, 1:3 **(H)**, 1:9 **(I)**. The significantly more abundant NK subsets are indicated with their *p*-values under each condition. The FDR is also indicated under each table. Bold green indicates an FDR<0.05 and a *p*-value <0.05. Unbold black indicates an FDR>0.05 still with a *p*-value <0.05.

### Long-term survival for BL patients treated with rituximab was lower for EBV-T2 tumors

To assess if EBV type influences patient outcomes, we performed Kaplan-Meier analysis for BL children treated at MTRH to avoid introducing variation between hospitals. We found that BL patients treated with rituximab (R-CC) had slightly better short-term survival (87.6%) compared to those treated with conventional chemotherapy alone (50%) (fig.8A, *p*=0.09). However, the long-term survival of rituximab-treated patients dropped to 57.6% one-year post diagnosis (fig.8B). For the R-CC treated BL patients, EBV type did not appear to impact short-term survival (fig.8C). Interestingly, the 2-year survival of EBV-T2-infected patients dropped to 62.5% while survival in those infected with EBV-T1 tended to stabilize at 71% (fig.8D). Since there are many factors that can impact long-term survival for pediatric cancer patients, this finding that points to more efficient ADCC for EBV-T1 would need further study to confirm.

**Fig. 8:**
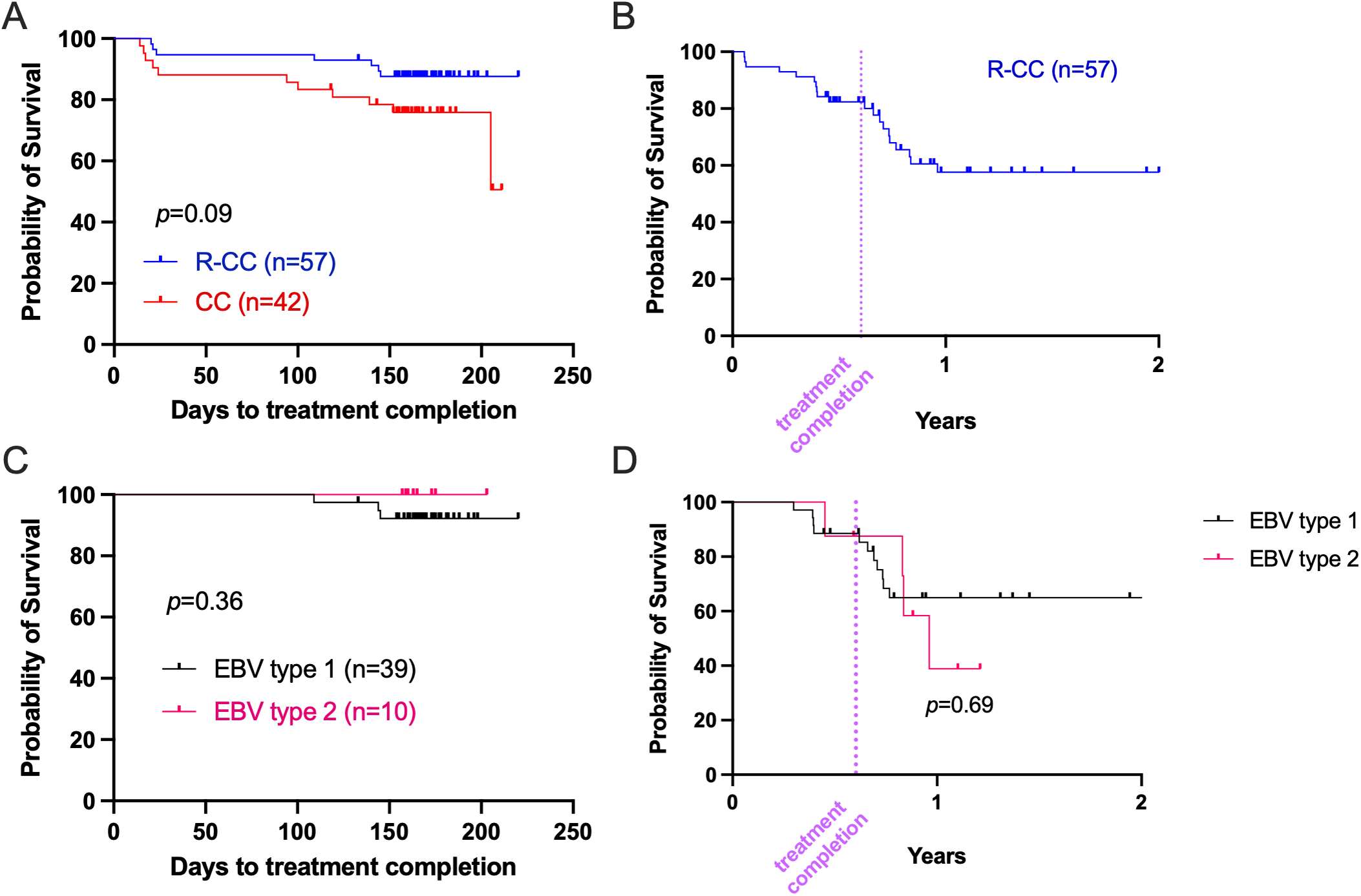
Rituximab confers short-term but not long-term survival advantage, which could be influenced by EBV type. **(A)** Kaplan-Meier analysis of BL survival to treatment completion between BL patients treated with (R-CC, in blue, n=57) or without (CC, in red, n=42) rituximab. **(B)** Kaplan-Meier curve of BL patients long term survival (in years) treated with rituximab (n=57). **(C)** Kaplan-Meier analysis of BL survival to R-CC treatment completion in regard to EBV-type (type 1 in black, n=33; type 2 in pink, n=7). **(D)** Kaplan-Meier analysis of R-CC-treated BL 2-year survival stratified by EBV-type (type 1 in black, n=33; type 2 in pink, n=7). All analysis excluded participants who died prior to the first course of treatment. Log-rank (Mantel-Cox) test was performed, and *p*-values are indicated on the graphs. R-CC: Rituximab and Combined Chemotherapy. CC: Combined Chemotherapy alone.

## Discussion

CD56^neg^ NK cells, historically overlooked due to their extremely low frequency in healthy individuals from more studied American and European populations, have gained increasing attention in the context of various chronic viral infections(5,36–38), in both pediatric and adult cancers(3,4,7,8). Despite their emerging relevance, their origin, functional role, and contribution to overall immune competence remain incompletely understood. In this study, we investigated the ADCC capacity of CD56^neg^ NK cells against EBV^pos^ BL cell lines using rituximab, a monoclonal antibody recognizing a quintessential B cell marker, CD20(9). Our findings demonstrate that although CD56^neg^ NK cells can mediate ADCC, they exhibit lower levels of degranulation compared to CD56^dim^ NK cells. This suggests that CD56^neg^ NK cells possess a subdued capacity to kill target cells mediated by ADCC, which appears to be complementary to the role of CD56^dim^ NK cells and not an alternative killing mechanism.

Since their initial description in the context of HIV(38), CD56^neg^ NK cells have been reported to exhibit poor direct cytotoxicity(3,5,39), likely due to the absence of CD56, N-CAM1 molecule, which is essential for immune synapse formation(40,41). However, CD56^neg^ NK cells exhibit transcriptomic and proteomic profiles strikingly similar to CD56^dim^ NK cells(4,7,42). CD56^neg^ NK cells express high levels of killer immunoglobulin-like receptors (KIRs), particularly KIR3DL1, along with CD57, NKG2C, NKG2D, perforin, and granzyme B, suggesting an adaptive or terminally differentiated NK cell phenotype. Despite having lower expression of NCRs such as NKp30, NKp46 and diminished direct cytotoxicity, CD56^neg^ NK cells from malaria-exposed individuals and pediatric BL patients do not exhibit an “exhaustion” signature with high and sustained expression of markers such as PD1, TIGIT, TIM3, and CTLA-4(4,43), indicating that they may still retain cytotoxic function. Moreover, Mavilio et. al demonstrated that CD56^neg^ NK cells could recover CD56 expression when cultured with recombinant IL-2 *in vitro*(44) suggesting that, even if these cells were on the path toward activation-induced cell death, their state is not beyond recovery. In this study, we observed a correlation between the abundance of CD56^neg^ NK cells and the levels of IgG1 and IgG3, the two subclasses known to mediate ADCC, and found that they could engage in ADCC against BL target cells but to a lesser degree compared to CD56^dim^ NK cells.

Our findings contrast those from a previous study examining malaria-exposed Ugandan children(43), which suggested that CD56^neg^ NK cells displayed greater ADCC activity compared to CD56^dim^ NK cells. This discrepancy may be attributed to several factors, including differences in experimental methodology and data interpretation. For instance, the authors only assessed the abundance of CD56^neg^CD107a^pos^ cells and did not evaluate the intensity of degranulation across NK subsets. Furthermore, they used plasma from malaria-exposed children to measure ADCC against *Pf*-infected red blood cells, whereas our study used rituximab to target BL cell tumors, suggesting that variation between NK ligands expressed by targets could explain this difference. Another important distinction is that their ADCC assay did not include TAPI-1, an inhibitor of ADAM17 metalloprotease which is known to shed CD16 after degranulation(45). Without CD16, CD56^neg^ NK cells cannot be properly isolated post-degranulation, as they merge with the non-NK cells (which are also CD56^neg^CD16^neg^). Moreover, despite performing NK cell enrichment, the authors did not report the purity of isolated NK cells or if it was assessed, leaving the possibility of contaminating non-NK cell populations (T-cells, granulocytes), which could also degranulate and skew the ADCC analysis. Given these methodological differences, it would have been ideal to sort CD56^neg^ NK cells prior performing the assay to accurately evaluate their contribution to ADCC. Unfortunately, this was not feasible in our study for several reasons, including the small blood volumes available from pediatric cancer patients, low number of total NK cells in their peripheral blood (∼5% compared to 10–15% in healthy adults), low yield of CD56^neg^ NK cells after sorting, and difficulty of expanding them in culture without converting them to a CD56^dim^ phenotype. Nevertheless, by assessing both the abundance of CD56^neg^CD107a^pos^ NK cells and the intensity of their degranulation, our results strongly suggest that, in response to rituximab against tumor cells, CD56^neg^ NK cells are capable of ADCC, albeit with reduced activity in comparison to CD56^dim^ NK cells.

Another noteworthy finding from our study is the differential response of NK cells to the different types of EBV within various BL cell lines. We observed significantly stronger direct cytotoxicity by NK cells against EBV-T2 BL tumors in contrast to EBV-T1. This effect is likely driven by differences in viral gene expression profiles between the two EBV types, as shown in our study. This finding is consistent with Shannon Kenney’s group using commercial BL cell lines(46). Our newly established EBV-T2 BL cell lines exhibited significantly higher expression of lytic genes, rendering them more susceptible to direct NK cell-mediated killing, particularly by the CD56^dim^ subset(47,48). Additionally, EBV-T2 BL cells expressed higher levels of IL-32, a cytokine known to enhance NK cell cytotoxic function(33). Notably, during acute EBV infection (infectious mononucleosis), an expansion of a CD56^dim^CD16^pos^ NKG2A^pos^KIR^neg^NKG2C^neg^CD57^neg^ NK cell subset has been described(47). These NK cells preferentially degranulate in response to EBV-infected B-cells expressing lytic antigens, reinforcing the idea that EBV-T2 tumors may be more vulnerable to direct NK cell detection. However, we did not observe differences in known NK ligand expression between EBV-T1 and -T2 BL cells, indicating other NK cell signaling mechanisms may be involved.

A limitation of this study was our inability to compare the level of supportive care over the study period and not having reliable 2-year survival information from children diagnosed at MTRH from March 2017 to June 2019. Nevertheless, we found that 1-year survival of R-CC-treated BL patients dropped to under 60%, close to survival rates of children not treated with rituximab. Rituximab has been shown to increase the risk of infection(49), including malaria in adult patients with lymphoma(50), which might be one of the causes of death of BL patients residing in malaria endemic areas. We also found that R-CC-treated BL patients infected with EBV-T2 tended to have lower long-term survival compared to those infected with EBV-T1. However, due to the small number of BL patients infected with EBV-T2 in our study population, this observation should be interpreted with caution. Altogether, our study raises the potential clinical relevance of determining EBV status in B cell tumors prior to initiating rituximab therapy.

To conclude, if a consequential proportion of the NK cell repertoire is skewed to be less cytotoxic this would have profound implications for viral immune surveillance and anti-cancer immunity as well as responsiveness to antibody-mediated therapies intended to engage NK cells. Therefore, a more personalized therapeutic approach should be considered to predict and optimize immuno-therapeutic options.

## Acknowledgements/Funding

The authors would like to thank the children and their families for participating in this study. This study was funded by the Alex Lemonade Stand Foundation (ALSF) Young Investigator Award (Forconi) and the NIH R01 NCI CA189806 (Moormann). Patient support was made available through a philanthropic donation from Takeda Pharmaceutical Company.

## Authorship contributions

A. C. S. F.: developed the hypothesis, designed/optimized all the assays and performed all the first set of cytotoxic assays, performed analysis and interpretations, wrote the manuscript.

B. L. S.: performed the second set of cytotoxic assays, reviewed the manuscript.

C. Z. R.: performed EBV qPCR, reviewed the manuscript.

V.M.B.: performed Malaria qPCR, reviewed the manuscript.

C.O.: performed the RNA-sequencing and analysis, reviewed the manuscript.

A. M.: performed the serological assays, reviewed the manuscript.

A. J. M.: performed the serological assays, reviewed the manuscript.

P.O.: performed computational analysis, reviewed the manuscript.

B.O.: collected all clinical data, reviewed the manuscript.

A. J. O: recruited participants, reviewed the manuscript.

F.N.: recruited participants, reviewed the manuscript.

T.A.V.: recruited participants, reviewed the manuscript.

A.K.: processing the participant samples, reviewed the manuscript.

J. B.: reviewed the RNA-sequencing analysis and interpretations, reviewed the manuscript.

C. M.: reviewed the cellular assays analysis and interpretations, reviewed the manuscript.

A. M. M.: reviewed all the analysis and interpretations, reviewed the manuscript.

