## Supplemental tables and figures for "EBV Type 1 versus Type 2: A determinant of NK cell anti-tumor activity in Burkitt lymphoma"

**Supp.table1: Reagents and Instruments information**

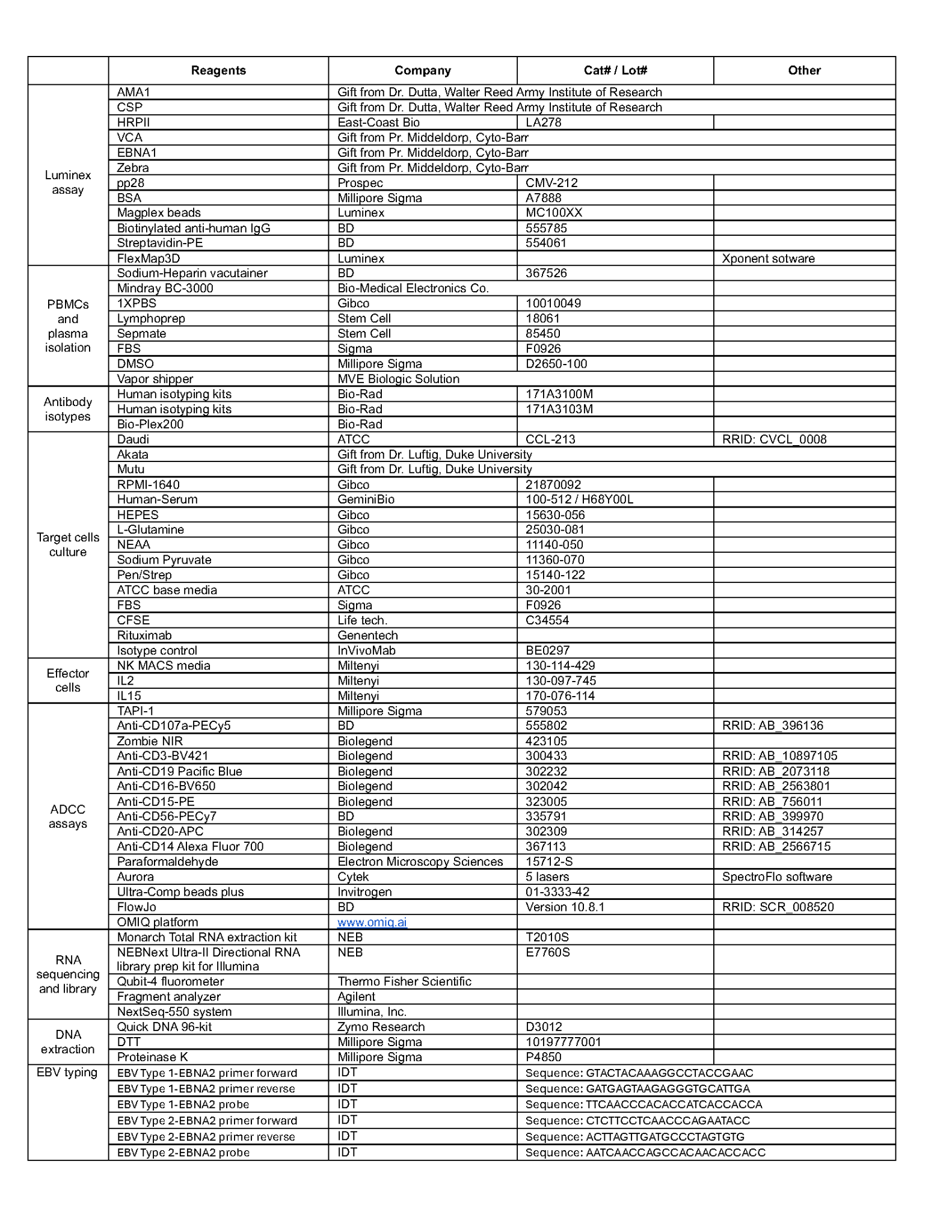

**Supp.table2: Population characteristics antibodies assays**

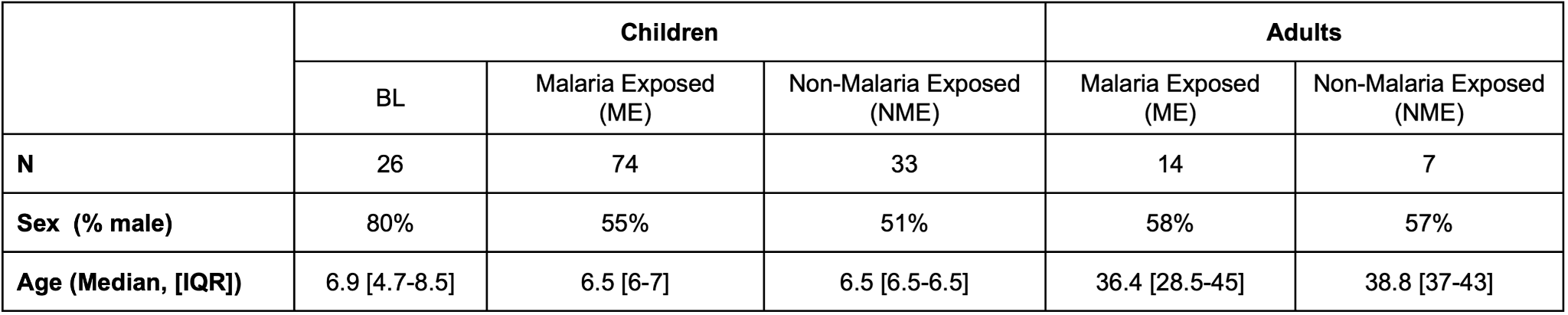

**Supp.table3: Population characteristics ADCC assays**

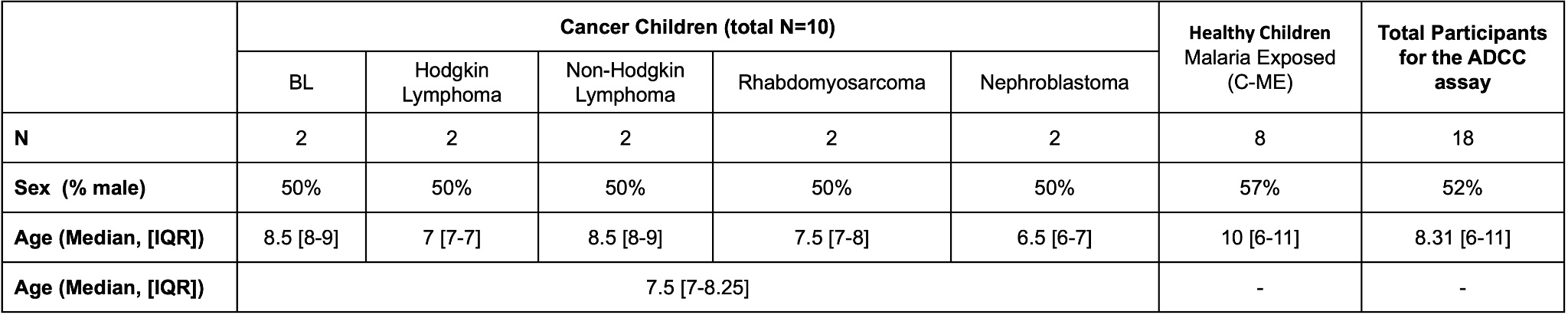

**Supp.table4: Population characteristics survival analysis**

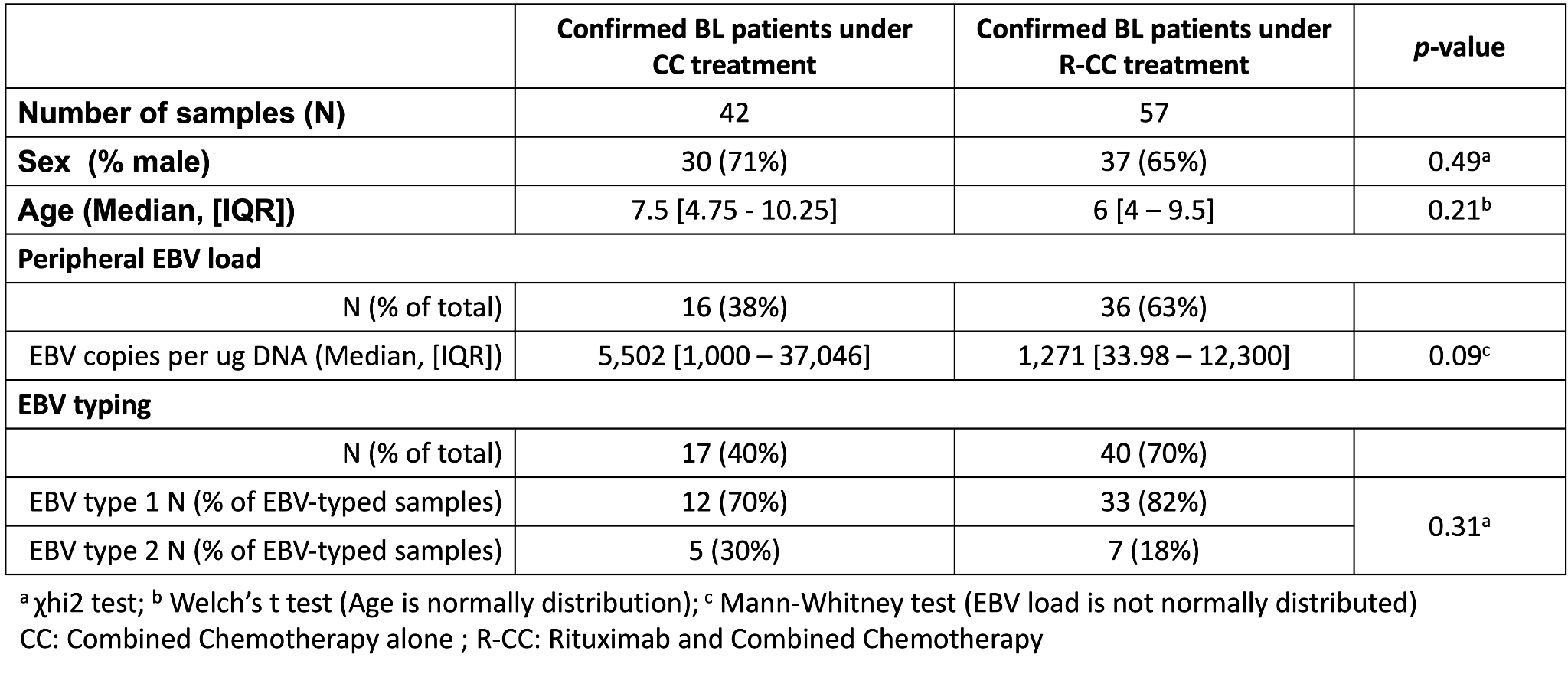

**Supp.table5: DEG analysis**

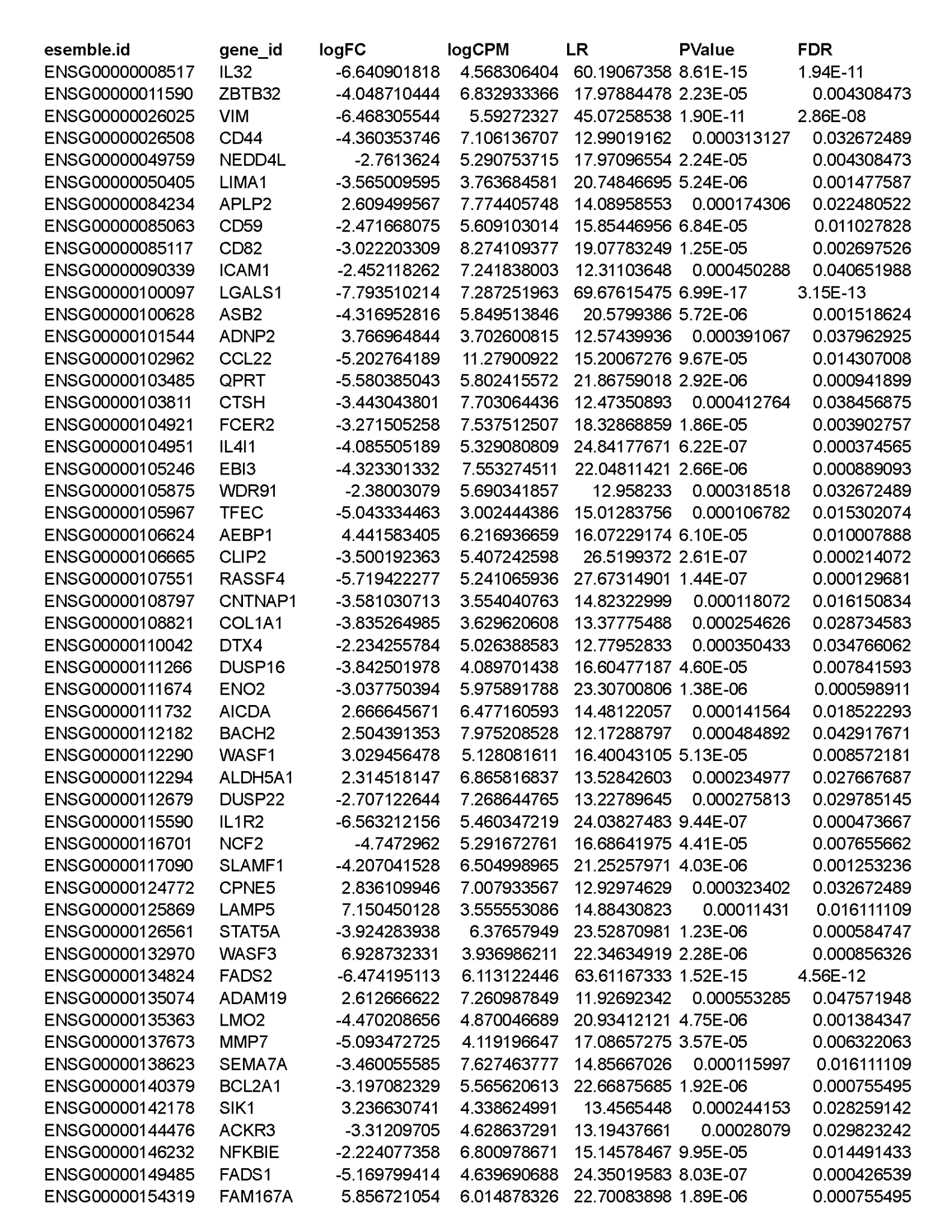

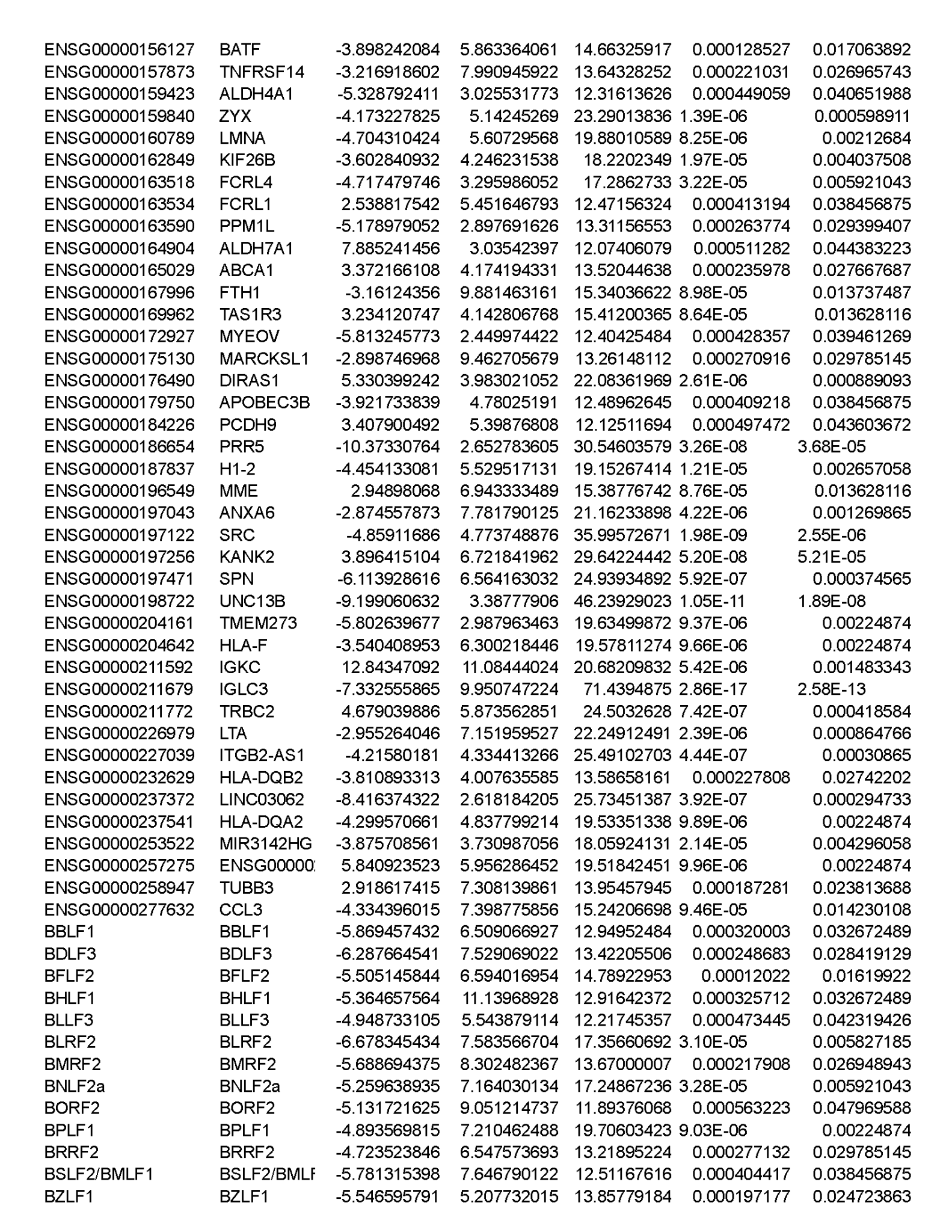

**Supp.Fig.1**

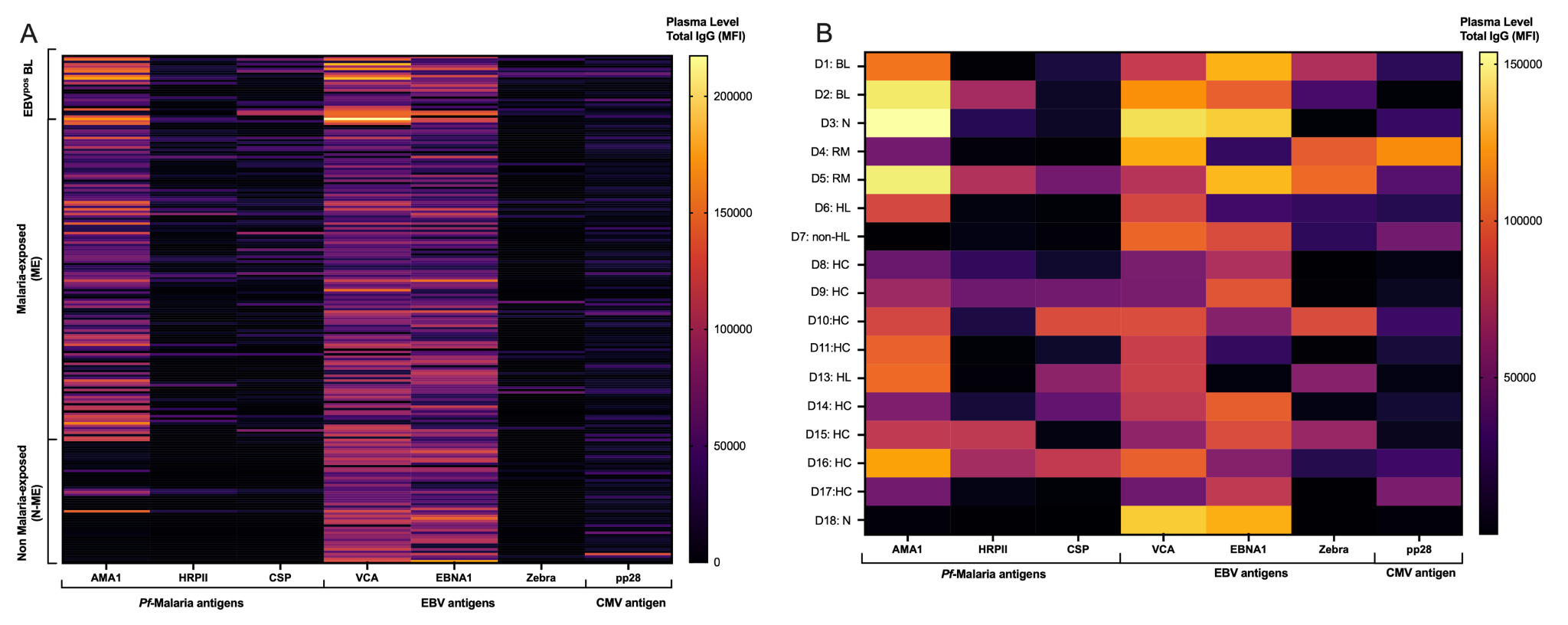

**Supp.Fig.1: Total IgG serological profile against malaria, EBV and CMV. (A)** Heatmap of total IgG antibodies against malaria (AMA1, HRPII, CSP), EBV (VCA, EBNA1, Zebra) and CMV (pp65) antigen. Participants were categorized into three groups: EBV-associated BL (EBV^pos^ BL), malaria-exposed (ME) and non malaria-exposed (N-ME). **(B)** Heatmap of total IgG antibodies against malaria, EBV and CMV from donors (D1 to D18) whom we used immune cells for the ADCC assay. The plasma from donor 12 was missing. For each donor, the clinical status is indicated: EBV-associated BL (BL), Nephroblastoma (N), Rhabdomyosarcoma (RM), Hodgkin-lymphoma (HL), non Hodgkin-lymphoma (non-HL), and healthy control (HC). The color scale indicates the median fluorescence intensity (MFI) of the total IgG plasma level for each participant. The yellow indicates high MFI whereas dark purple/black indicates low MFI.

**Supp.Fig.2**

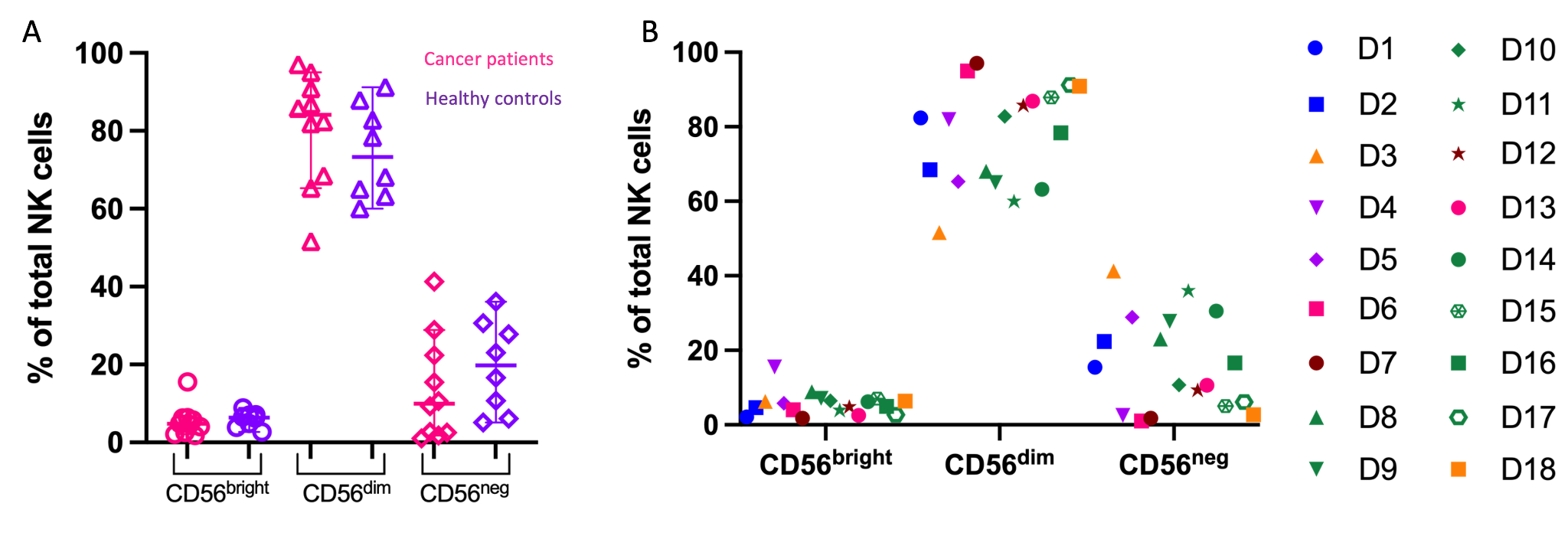

**Supp.Fig.2: NK cell main subsets distribution.** Abundance of CD56^bright^, CD56^dim^ and CD56^neg^ NK cells subsets within total NK cells shown by clinical status of the participants **(A)**: participants with cancer (pink), healthy controls (purple); and shown by participants D1 to D18 **(B)**. Participants with EBV^pos^ BL are in blue, Nephroblastoma in orange, Rhabdomyosarcoma in purple, Hodgkin-lymphoma in pink, non Hodgkin-lymphoma in brown, and healthy control in green.

**Supp.Fig.3**

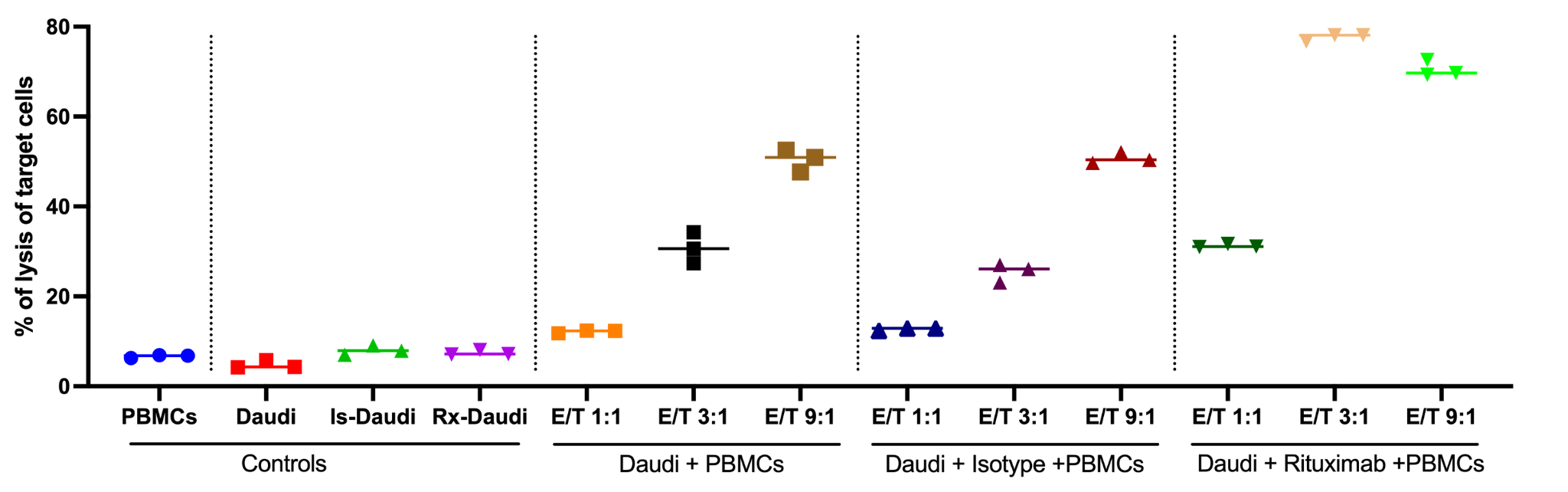

**Supp.Fig.3: Validation of the ADCC assay.** PBMCs from a healthy Kenyan adult were used in triplicate to perform ADCC against opsonized Daudi cell lines.

**Supp.Fig.4**

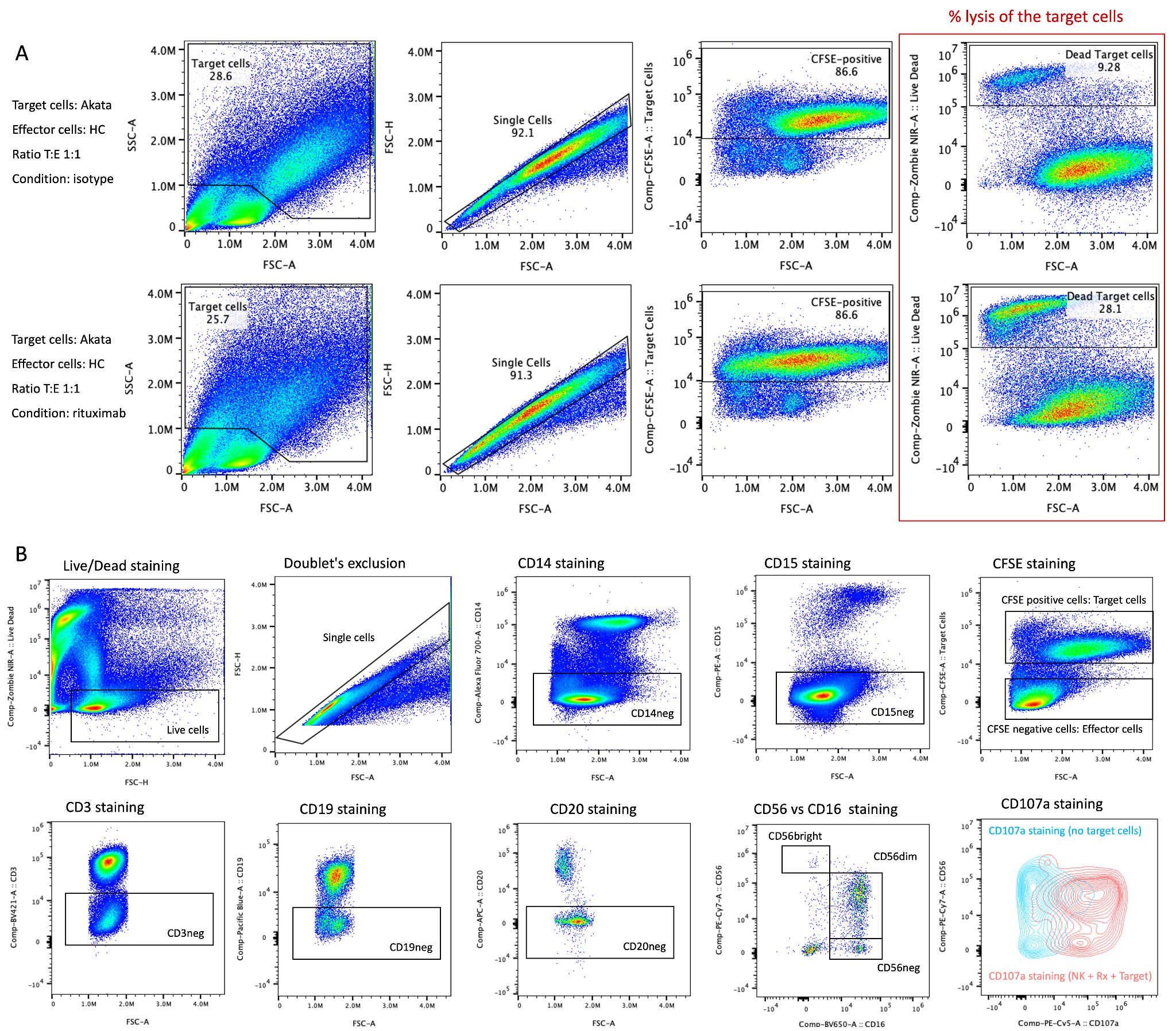

**Supp.Fig.4: Flow cytometry gating strategy assessing the percentage of lysis of the target cells (A) as well as the degranulation of the effector cells (B).**

**Supp.Fig.5**

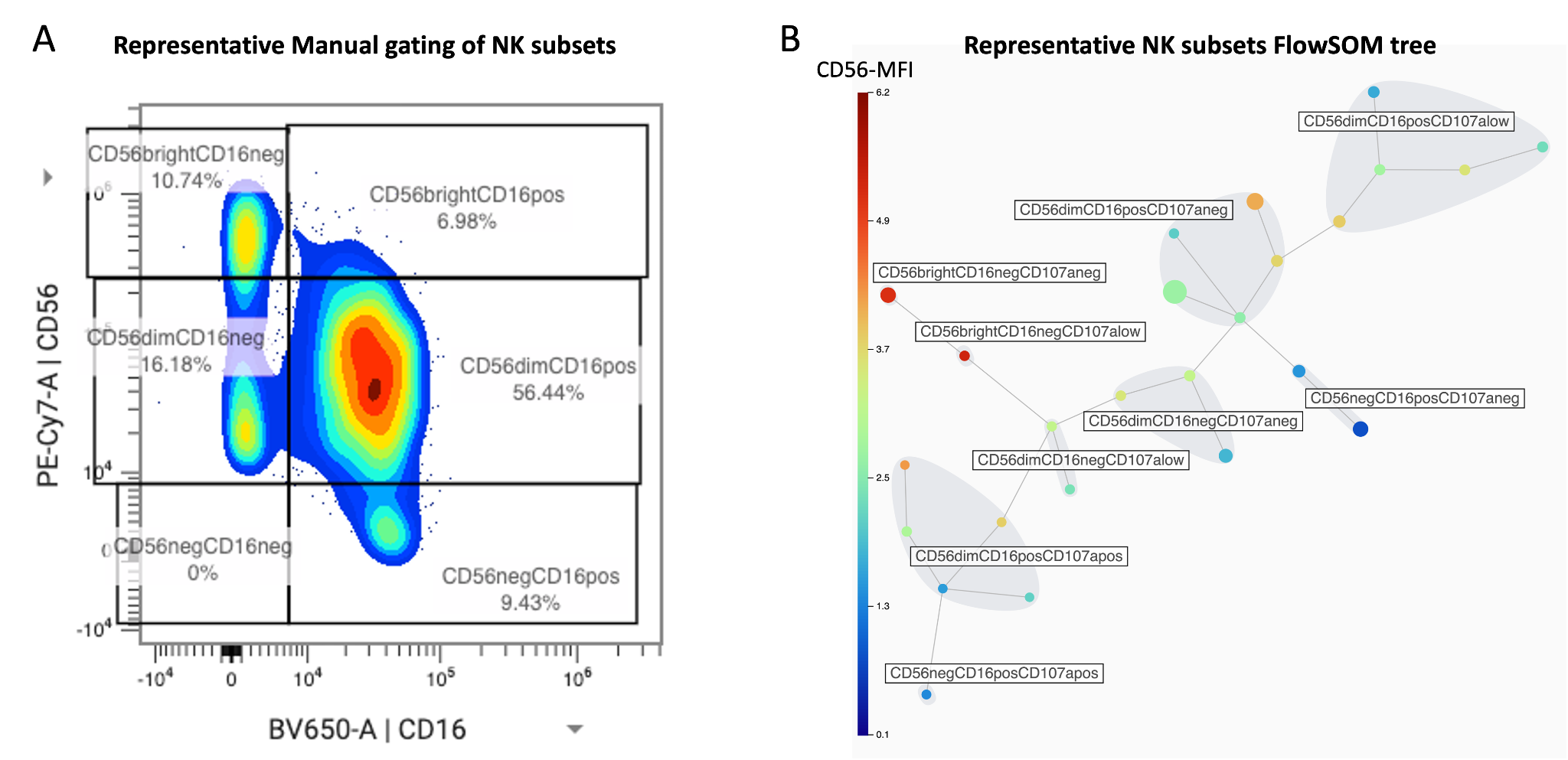

**Supp.Fig.5: NK subsets identified by manual gating (A) versus FlowSOM clustering analysis (B).**

**Supp.Fig.6**

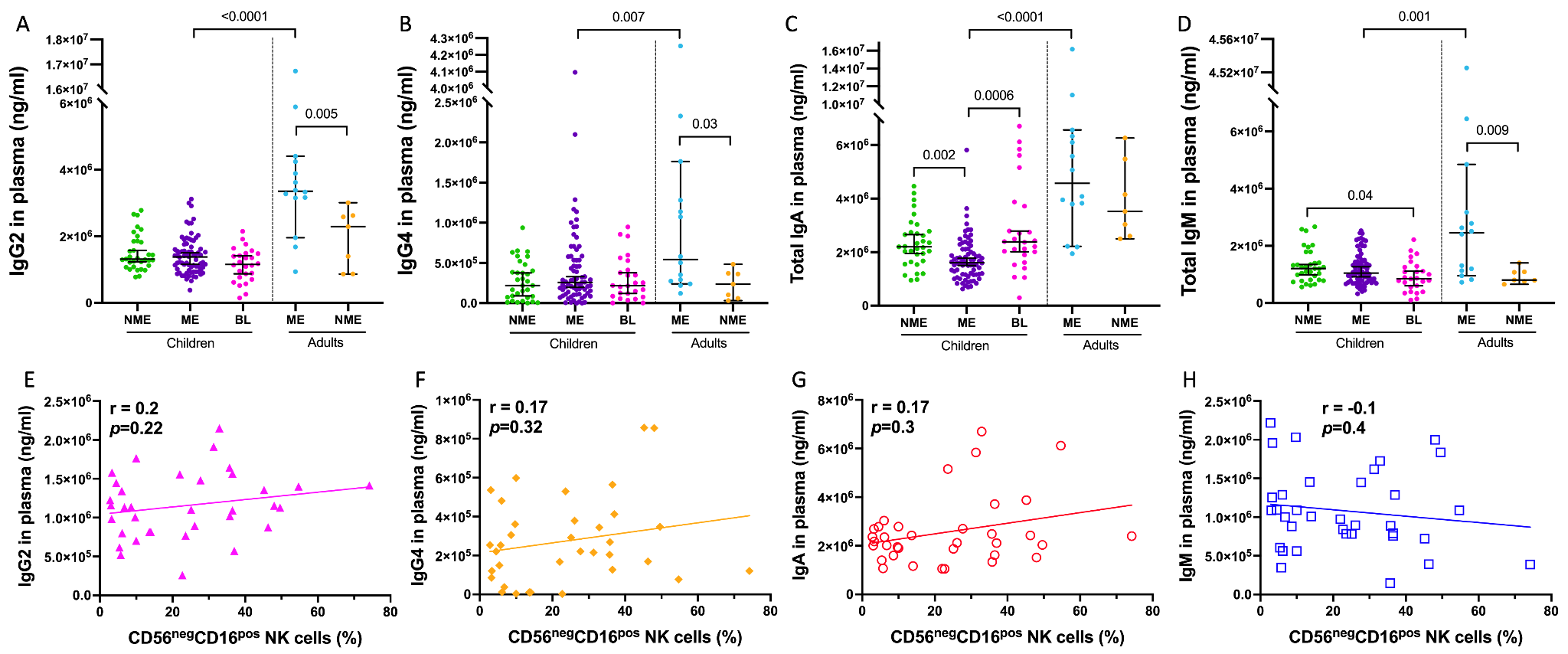

**Supp.Fig.6: No correlation between plasma levels of IgG2, IgG4, total IgA and total IgM with the abundance of peripheral CD56^neg^ NK cells.** Dot plots of plasma level of IgG2 **(A)**, IgG4 **(B)**, total IgA **(C)** and total IgM **(D)** from non-malaria exposed (NME, n=33, in green) and malaria exposed (ME, n=74, in purple) children, BL patients (BL, n=26, in pink), ME and NME adults (n=14, blue and n=7, orange, respectively). Median and 95% confidence intervals are represented, significant *p*-values found comparing the children groups with the Kruskal-Wallis test are indicated on the plot while significant *p*-values comparing the two adults groups or the two ME groups are indicated following Mann-Whitney test. Correlation test assessing relationships between plasma levels of IgG2 **(E)**, IgG4 **(F)**, total IgA **(G)** and total IgM **(H)**, and the frequency of peripheral CD56^neg^CD16^pos^ NK cells from total NK cells from all children participants. Plasma levels of IgG2, IgG4, IgA and IgM did not follow a normal distribution, therefore, the Spearman test was used (r and *p*-values indicated on the plots).

**Supp.Fig.7**

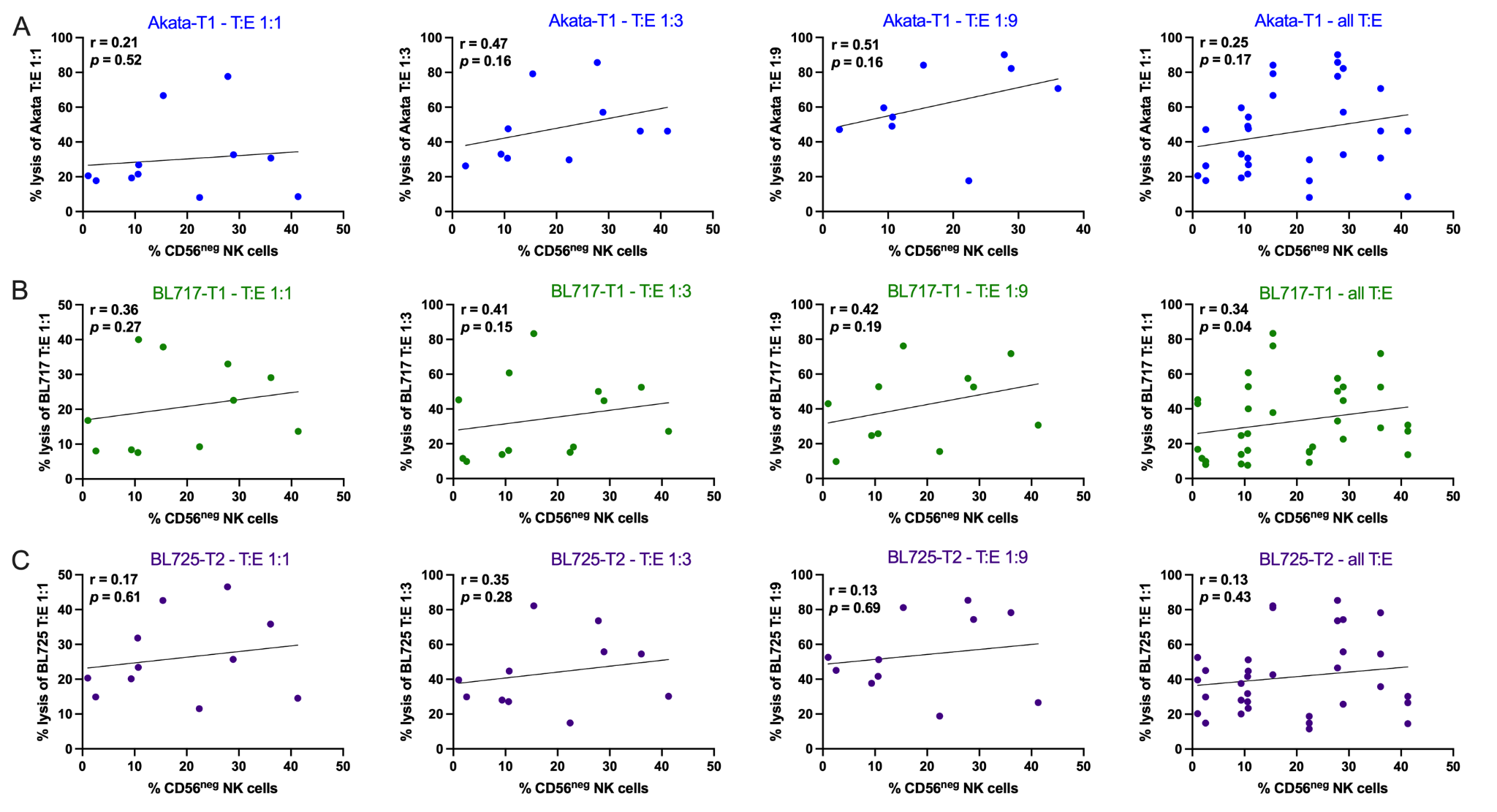

**Supp.Fig.7: No correlation between the percentage of lysis of the rituximab-treated target cells and the percentage of CD56^neg^ NK cells.** Because of the non-normal distribution of the data, the Spearman test was used to assess relationships between the percentages of CD56^neg^ NK cells and the percentage lysis of the target cells Akata **(A)**, BL717 **(B)** and B:725 **(C)** at different T:E ratios (r and *p*-values indicated on the plots).

**Supp.Fig.8**

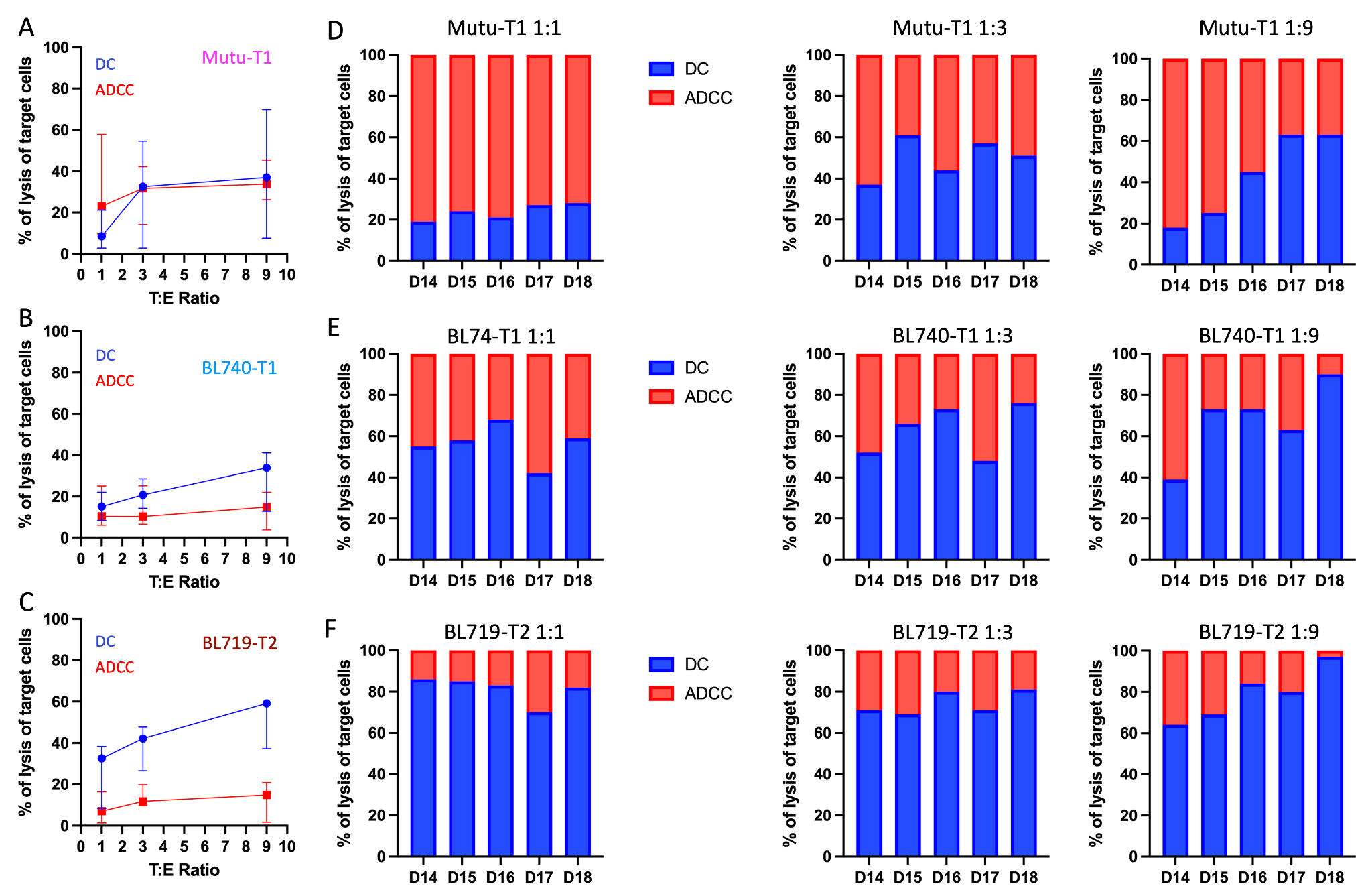

**Supp.Fig.8: ADCC magnitude against Mutu, BL740 and BL719.** Range of percentage of lysis of Mutu **(A)**, BL740 **(B)**, and BL719 **(C)** by DC (in blue) or ADCC (in red) across three different T:E ratios. The symbol represents the median and the bar the error with 95% confidence interval. The proportions of DC (in blue) and ADCC (in red) killing of Mutu **(D)**, BL740 **(E)**, and BL719 **(F)** are then reported with bar plots for each participant (D14 to D18), for the three different T:E ratios.

**Supp.Fig.9**

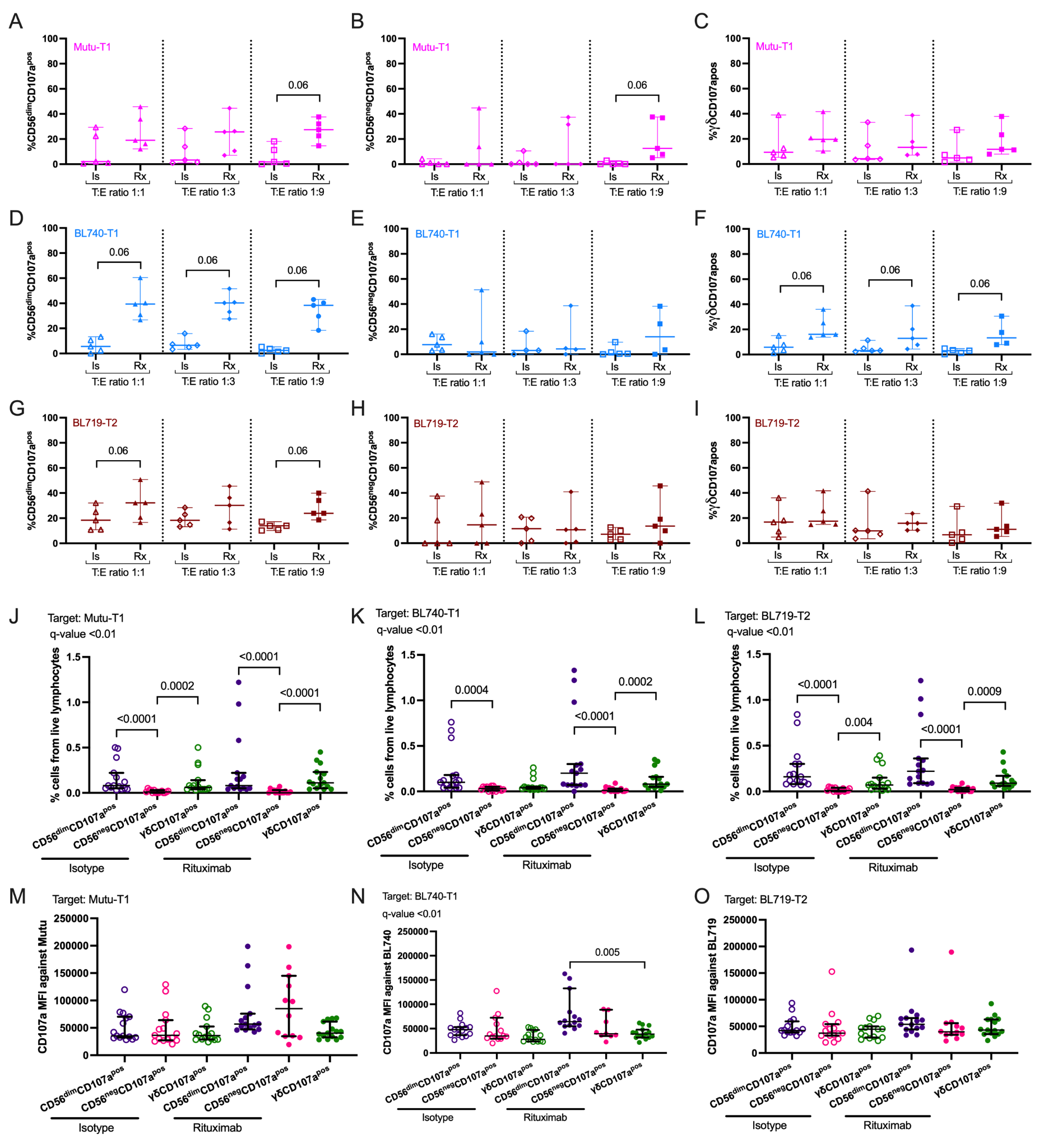

**Supp.Fig.9: CD56^dim^ NK cells degranulate more in presence of rituximab against EBV-T1 BL cell lines.** Percentage of CD56^dim^CD107a^pos^ cells from total CD56^dim^ NK cells **(A)**, percentage of CD56^neg^CD107a^pos^ cells from total CD56^neg^ NK cells **(B)**, and percentage of γδ^pos^CD107a^pos^ cells from total γδ T-cells after co-culture with Mutu, BL740 (D, E, and F, respectively) and BL719 (G, H, and I, respectively), in presence of isotype (empty symbols) or rituximab (full symbol). All the data are represented with median and 95% confidence intervals. Two-tailed Wilcoxon paired test was used to compare the isotype condition to the rituximab treatment. Percentage of CD56^dim^CD107a^pos^ (purple), CD56^neg^CD107a^pos^ (pink), and γδ^pos^CD107a^pos^ (green) cells from total live lymphocytes after co-culture with Mutu **(J)**, BL740 **(K)** and BL719 **(L)**, in presence of isotype (empty symbols) or rituximab (full symbol). CD107a median fluorescence intensity (MFI) of CD56^dim^CD107a^pos^ (purple), CD56^neg^CD107a^pos^ (pink), and γδ^pos^CD107a^pos^ (green) cells after co-culture with Mutu **(M)**, BL740 **(N)** and BL719 **(N)**, in presence of isotype (empty symbols) or rituximab (full symbol). Data are represented with median and 95% confidence intervals. Multiple comparisons were performed using Kruskal-Wallis statistical test with FDR correction. Both *p*-values and *q*-values (adjusted *p*-values based on the FDR) are indicated on the plots.

**Supp.Fig.10**

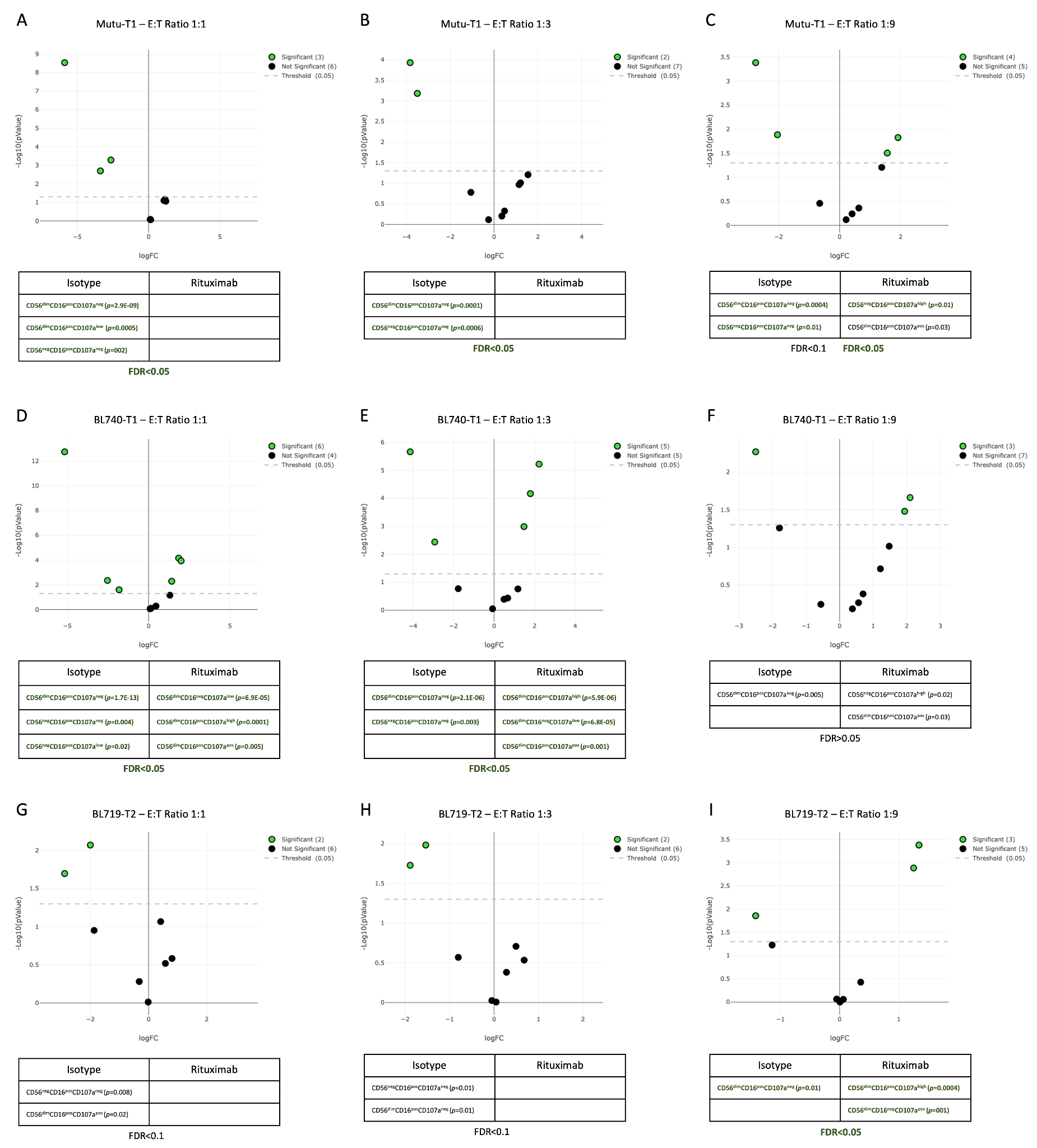

**Supp.Fig.10:** EdgeR analysis following a semi-supervised FlowSOM clustering showing, in green, the NK subsets statistically differently abundant between the isotype and the rituximab condition after co-culture with Mutu T:E ratio 1:1 **(A)**, 1:3 **(B)**, 1:9 **(C)**; BL740 T:E ratio 1:1 **(D)**, 1:3 **(E)**, 1:9 **(F)**; and BL719 T:E ratio 1:1 **(G)**, 1:3 **(H)**, 1:9 **(I)**. The significantly more abundant NK subsets are indicated with their *p*-values under each condition. The FDR is also indicated under each table. Bold green indicates an FDR<0.05 and a *p*-value <0.05. Unbold black indicates an FDR>0.05 still with a *p*-value <0.05.
